# Structure of YbbAP-TesA: a Type VII ABC transporter–lipid hydrolase complex

**DOI:** 10.1101/2025.05.16.654284

**Authors:** Martin B.L. McAndrew, Jonathan Cook, Amy Gill, Kavya Sahoo, Clare Thomas, Phillip J. Stansfeld, Allister Crow

**Affiliations:** School of Life Sciences, University of Warwick, United Kingdom; Warwick Medical School, University of Warwick, United Kingdom; Department of Chemistry University of Warwick, United Kingdom

**Keywords:** Type VII ABC Transporters, Lysophospholipase, Structural Microbiology, Bacterial Cell Envelope

## Abstract

Type VII ABC transporters are ATP-powered membrane protein complexes that drive key biological processes in the bacterial cell envelope. In *E. coli*, three of the four Type VII ABC systems have been extensively characterised including: the FtsEX-EnvC cell division complex, the LolCDE-LolA lipoprotein trafficking machinery and the MacAB-TolC efflux pump. Here we describe a fourth *E. coli* Type VII ABC system, YbbAP-TesA, which combines a Type VII ABC transporter with a multifunctional hydrolytic enzyme. Structures of the complete YbbAP-TesA complex, and of YbbAP with and without bound ATP analogues, capture the long-range transmembrane conformational changes that are the hallmark of this ABC superfamily’s mechanotransmission mechanism. We further show that YbbAP-TesA can hydrolyse a variety of ester and thioester substrates and experimentally confirm a constellation of active site residues in TesA. Our data suggests YbbAP has a role in extracting hydrophobic molecules from the inner membrane and presenting these to TesA for hydrolysis. The work extends collective knowledge of the remarkable diversity of the ABC superfamily and establishes a new function for Type VII ABC transporters in bacterial cells.

**Significance Statement:** Type VII ABC transporters are unique to bacteria and play important roles in bacterial physiology including cell division, antibiotic resistance, siderophore secretion, antibiotic sensing, toxin secretion, biofilm regulation and lipoprotein trafficking. Here we identify a complex that combines an atypical Type VII ABC transporter (YbbAP) with a periplasmic lipid hydrolase (TesA). The YbbAP-TesA complex is structurally and functionally distinct from all known Type VII ABC transporter systems including FtsEX-EnvC, LolCDE-LolA, BceAB-BceS and MacAB-TolC. Structures suggest that YbbAP-TesA uses ATP-driven mechanotransmission to extract substrates from the inner membrane and hydrolyse them in the periplasmic space. The discovery and characterisation of YbbAP-TesA highlights the structural and functional diversity of Type VII ABC transporter complexes and suggests a new function for these proteins in the bacterial cell envelope.

## Introduction

ABC transporters belong to multiple superfamilies that are recognised by the 3D fold of their transmembrane domain and the presence of a cytoplasmic ATP-Binding Cassette (1). In *E. coli*, there are four Type VII ABC transporters: MacB, LolCDE, FtsEX and YbbAP (2). MacB forms the inner membrane component of the MacAB-TolC tripartite efflux pump which provides resistance to macrolide antibiotics and drives secretion of small protein toxins from the periplasm to the extracellular space (3–5). LolCDE couples the ATP binding and hydrolysis cycle to the extraction of lipoproteins from the inner membrane and passes them to the periplasmic chaperone (LolA) for delivery to the outer membrane (6–11). FtsEX-EnvC allosterically regulates periplasmic amidases that cleave the peptidoglycan layer during cell division (12–21). The function of YbbAP is heretofore unknown, but its predicted topology differs from the other *E. coli* Type VII ABC systems due to the presence of 10 transmembrane helices rather than the four that is typical of this protein family (2, 22).

The Type VII ABC transporters all share a conserved transmembrane fold and operate via a mechanotransmission mechanism in which the ATP-binding and hydrolysis cycle is coupled to transmembrane conformational changes that drive work on the periplasmic side of the bacterial inner membrane (1–3). Typically, and unlike most conventional ABC transporters, such work does not involve the passage of substrates across the cytoplasmic membrane. Instead, Type VII ABC transporters drive a variety of energetic processes in the periplasm (or extra-cytoplasmic space) including extraction of molecules from within the bacterial membrane or activation of periplasmic enzymes.

A recurring theme within the Type VII ABC superfamily is the association of the ABC transporter component with a partner protein (or proteins) that define the complex’s biological function. For example, several Type VII ABC transporters form tripartite efflux pumps through interactions with periplasmic adaptor proteins and TolC-like outer membrane proteins. The best studied case is MacAB-TolC (3, 4), although many similar systems can be identified such as the aap/dispersin-secreting systems (23, 24) and the pyoverdine secreting PvdT system found in *Pseudomonas aeruginosa* (25). Other non-efflux pump examples include the FtsEX-like protein systems that control the hydrolysis of the peptidoglycan layer via conformational changes in a periplasmic activator (EnvC) (20, 14, 15, 13, 18) or the extraction of lipoproteins by LolCDE for delivery to the outer membrane via chaperones such as LolA (6–10, 26–29). A subset of Type VII ABC transporters also interact with partners located in the membrane – for example, the BceAB-like systems involved in bacitracin resistance interact with a dimeric transmembrane histidine kinase (BceS) involved in antibiotic detection and intracellular signalling (30–33). Identification of partner proteins can therefore be used to both classify and predict the functions of uncharacterised Type VII ABC transporters and to identify novel systems that are yet to be characterised. Recently, a revolution in computational protein structure prediction has facilitated the prediction of both individual protein structures and larger multi-protein assemblies (34–36). In silico ‘co-folding’ of proteins alongside libraries of potential partners holds great promise for identification of functionally distinct Type VII ABC transporters – especially where such predictions can be verified by experiment.

Here we identify and examine an intriguing partnership between a Type VII ABC transporter (YbbAP) and a periplasmic esterase (TesA). TesA is a periplasmic lysophospholipase with both esterase and thioesterase activity (37, 38). Our data suggests that YbbAP-TesA is a novel Type VII ABC transporter system that extracts hydrophobic compounds from the bacterial inner membrane and presents them for hydrolysis by TesA in the periplasm.

## Results

### TesA is the periplasmic partner of the YbbAP ABC transporter

YbbAP is the only Type VII ABC transporter from *E. coli* that is yet to be structurally and functionally characterised (2). To identify potential partners of YbbAP that define its function, we performed a brute-force ‘*in silico* pulldown’ using AlphaFold. The YbbP protein was independently co-folded alongside all 534 predicted periplasmic proteins from *E. coli* and the complexes scored according to measures of prediction confidence (pDockQ) and structural plausibility (Molprobity clashscore) (34, 39). This approach identified TesA as a strong candidate for a periplasmic partner of YbbAP (**Fig. 1a**). TesA is a multifunctional periplasmic enzyme belonging to the GDSL family of serine esterases and lipases and is widely annotated as a Lysophospholipase (37, 38, 40). The interaction between YbbAP and TesA hydrolase is distinct from the other Type VII ABC systems that have been characterised suggesting a novel biological function for YbbAP-TesA. We therefore sought to further characterise the structure and function of YbbAP-TesA.

**Figure 1:**
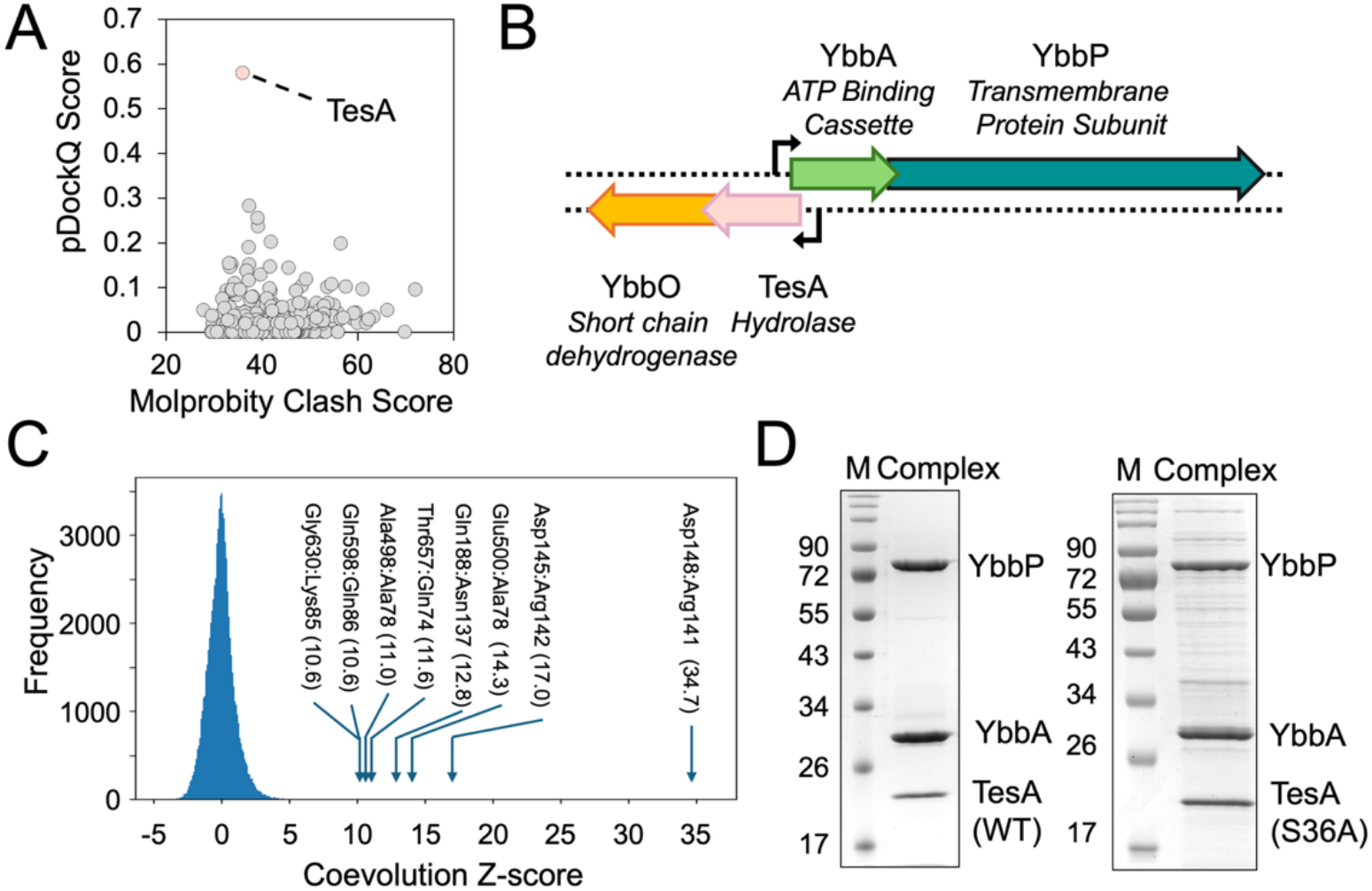
Identification of TesA as the periplasmic partner of YbbAP. (a) Pairwise *in silico* co-folding of YbbP with all sec/tat secreted proteins from *E. coli*. Each co-fold attempt is represented on the 2D plot by their pDockQ score and Molprobity clash score. The TesA-YbbP pair is indicated. (b) Genomic arrangement of YbbA, YbbP, TesA and YbbO genes in *E. coli*. (c). Coevolution analysis between YbbP and TesA. All inter-protein residues were scored for coevolution using CCMPred, converted to a Z-score and depicted as a frequency histogram. Coevolving residue pairs between YbbP and TesA with Z-scores above 10 are annotated (Further co-evolving residue pairs within the YbbAP-TesA complex are given in **Table S1**). (d) SDS-PAGE gel showing co-purification of YbbAP-TesA as a complex.

### YbbAP and TesA are co-located in the *E. coli* genome

Additional support for the interaction between YbbP and TesA was provided by its genetic context. We inspected the location of YbbA, YbbP and TesA encoding genes using KEGG (41). The *tesA* gene is located upstream of *ybbA* and *ybbP* in the *E. coli* genome and shares an operon with *ybbO* (annotated as a putative short chain dehydrogenase or aldehyde reductase). The putative *tesA*-*ybbO* operon partially overlaps the neighbouring *ybbA-ybbP* operon which faces the opposite direction on the chromosome (**Fig. 1b**). Similar genetic arrangements uniting TesA with YbbAP are found among other Gram-negative bacteria including *Pseudomonas aeruginosa* for which TesA has also been characterised as a lysophospholipase (38). The co-localisation of YbbAP and TesA in the genome strongly suggests these proteins may be functionally related and possibly co-regulated.

### Coevolution analysis predicts a direct interaction between YbbP and TesA

To further test the likelihood of the transporter-hydrolase interaction we identified co-evolving residue pairs between YbbP and TesA. Using CCMPred (42), we identified several high-scoring residue pairs at the interface between YbbP and TesA, and between YbbA and YbbP, suggesting intermolecular contacts (**Fig. 1c** and **Table S1**). As a control, we applied the same co-evolutionary contact analysis to TesA and YbbA which are located on opposite sides of the cytoplasmic membrane. As expected, we found no evidence of a signal between YbbA and TesA further strengthening our confidence in the strong coevolutionary signals we found between YbbA and YbbP, and between YbbP and TesA.

### Co-purification of YbbAP-TesA experimentally confirms the complex

We next experimentally tested whether YbbAP-TesA forms a physical complex by co-purifying YbbAP and YbbAP-TesA. We over-expressed TesA alongside the His-tagged YbbAP ABC transporter and successfully purified a three-protein complex in detergent using Ni-Immobilised Metal Affinity Chromatography (**Fig. 1d**). Addition of ATP (or an ATP analogue) did not affect the co-purification of YbbAP with TesA and neither did introduction of a catalytic site mutation (Ser36Ala) in TesA. These results confirm that YbbAP forms a stable complex with TesA and that YbbAP-TesA represents a novel membrane protein complex in which a periplasmic lysophospholipase is bound to the Type VII ABC transporter.

### CryoEM structures of YbbAP and the YbbAP-TesA complex

We next determined structures of YbbAP and the YbbAP-TesA complex using cryoEM. Statistics for each structure determination are given in **Table S2** and figures showing the quality of the underpinning cryoEM maps appear in **Figs. 2a** and **2b**. Workflow diagrams for cryoEM processing are shown in **Figs. S1, S2, S3**. The best resolved structure is that of YbbAP bound to an ATP analogue which we determined at 3.7 Å resolution (**Fig. 2a**). Structures of nucleotide-free YbbAP and the complete YbbAP-TesA complex (including a Ser36Ala catalytic site mutation in TesA) were determined at 4.0 Å and 4.6 Å resolution, respectively. All three structures have well-defined protein cores but exhibit greater mobility at their periphery. Secondary structure elements, transmembrane helices and the periplasmic subdomains are all well-resolved as are the boundaries of the detergent micelles that support the transmembrane domain. The key structure of YbbAP-TesA is underpinned by a high quality cryoEM map that unambiguously positions TesA relative to YbbAP and resolves mechanistically important features such as the YbbAP-TesA interface and the bound nucleotide analogues (**Fig. 2b**).

**Figure 2:**
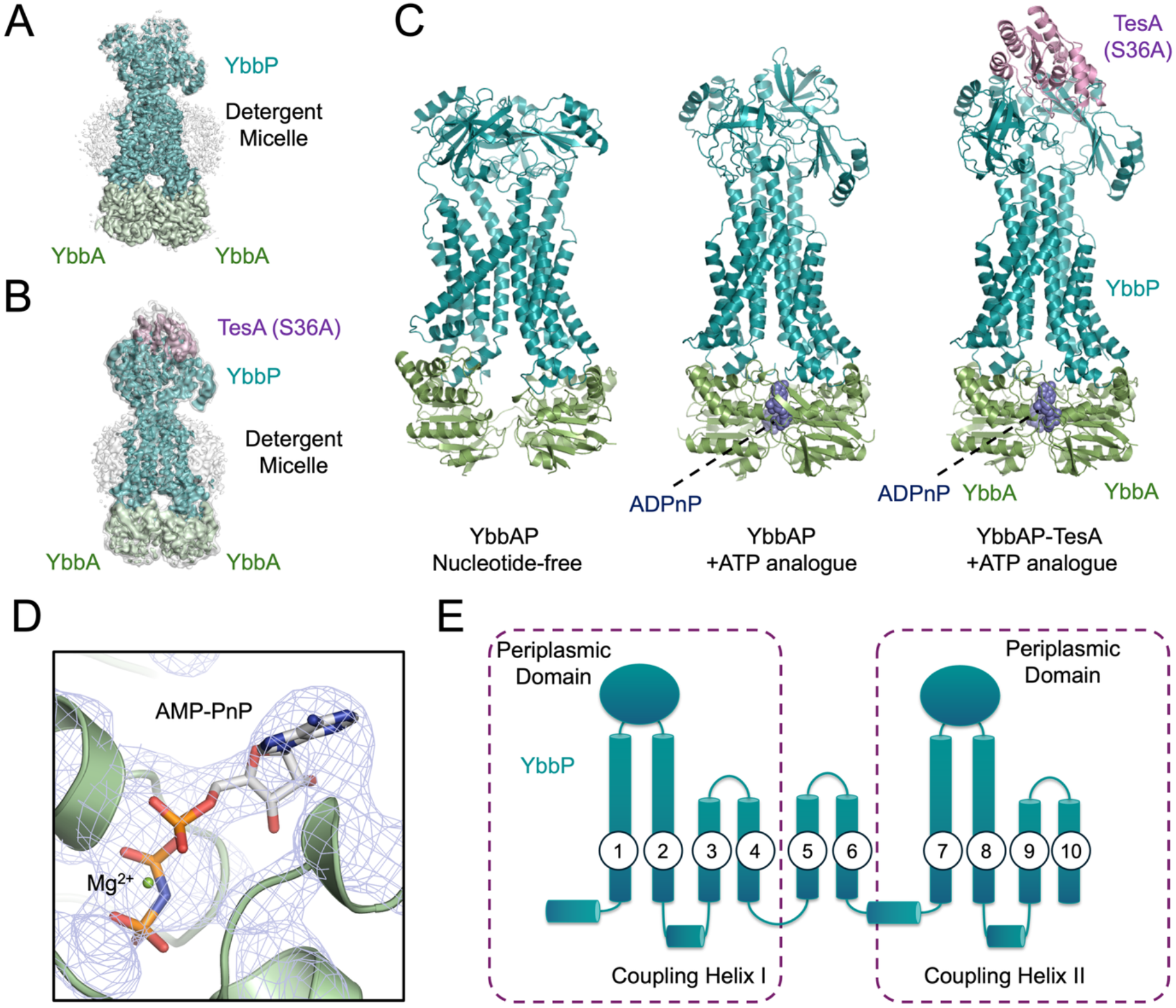
Structures of YbbAP and the YbbAP-TesA complex. (a) CryoEM map for the ATP-bound YbbAP at 3.6 Å resolution. (b) CryoEM map for ATP-bound YbbAP-TesA complex contoured to visualise the detergent belt. Maps are shown at two contour levels – a partially transparent low contour map is shown in *white* (4.5 σ) and a high contour map is shown coloured by nearest protein chain (7 σ). (c) Structures of apo YbbAP, ATP analogue-bound YbbAP, and ATP analogue-bound YbbAP-TesA. Proteins are shown in cartoon representation (YbbA *green*, YbbP *teal* and TesA *pink*) and the non-hydrolysable ATP analogue (AMP-PNP) is shown in *blue* atomic spheres. (d) ATP binding site in the YbbAP-TesA structure. The blue mesh shows the cryoEM map contoured at 8 σ. AMP-PNP is shown in sticks and a Mg^2+^ ion as a *green* sphere. (e) Topology diagram for YbbP. Dashed boxes indicate the duplicated segments that give YbbP its internal pseudosymmetry.

The three protein structures are shown side-by-side in **Fig. 2c**. The complete YbbAP-TesA complex has a 2:1:1 stoichiometry with two molecules of the ATP binding cassette (YbbA) bound to the cytoplasmic side of YbbP, and a single molecule of TesA bound on the periplasmic side of YbbP. YbbAP-TesA has 10 transmembrane helices and a similar transmembrane topology to the BceABS family of Type VII ABC transporters (31, 32). The ATP analogue used to stabilise the ATP-bound conformation is well resolved in both the YbbAP-TesA and YbbAP structures and binds in the conventional manner, sandwiched between the two ABC domains (**Fig. 2d**).

A diagram of the YbbP topology is shown in **Fig. 2e**. A large periplasmic domain is present between transmembrane helix 1 and 2, and between transmembrane helix 7 and 8. The YbbP periplasmic domains are similar in fold to the periplasmic domains of MacB and LolCDE and contain both the Porter and Sabre subdomains. As with other Type VII ABC transporters, the periplasmic domains are raised above the plane of the membrane by helical extensions of the connecting transmembrane segments. In the ATP-bound structures, these four helices form a tight 4-helix bundle, as is also seen in ATP-bound conformations or MacB (3) and LolCDE (8, 10, 11, 26). On the cytoplasmic side of the membrane, YbbP interacts with each YbbA monomer via two separate coupling helices. The first coupling helix of YbbP is located between transmembrane helix 2 and 3 and the second between transmembrane helix 8 and 9. The YbbP fold is pseudosymmetric due to an ancient duplication of the first and last four transmembrane segments. The internal symmetry of YbbP therefore mimics the dimeric structure seen for MacB and LolCDE with an additional pair of transmembrane helices (TM 5 and 6) serving as a link between the duplicated N- and C-terminal sequences (**Fig. 2e**). Overall, the three structures show significant conformational differences related to the nucleotide status of the ATP binding cassettes. The structural differences between ATP-bound and nucleotide free YbbAP span the entire protein complex and imply conformational changes on both the cytoplasmic and periplasmic sides of the membrane.

### Evidence for a mechanotransmission mechanism in YbbAP-TesA

Type VII ABC transporters undergo long-range transmembrane conformational changes that are intrinsically tied to their biological function and driven by ATP binding and hydrolysis. Such conformational changes have been termed mechanotransmission since it is ultimately mechanical forces that are propagated across the membrane rather than substrates (2, 3). We compared the structures of YbbAP solved with and without a non-hydrolysable ATP analogue to assess whether YbbAP-TesA also uses a mechanotransmission mechanism. A movie showing a molecular morph between the nucleotide-free and ATP-bound states is given as supplemental information (**Movie 1**) and a comparison of the structures is shown in **Figure 3**.

**Figure 3:**
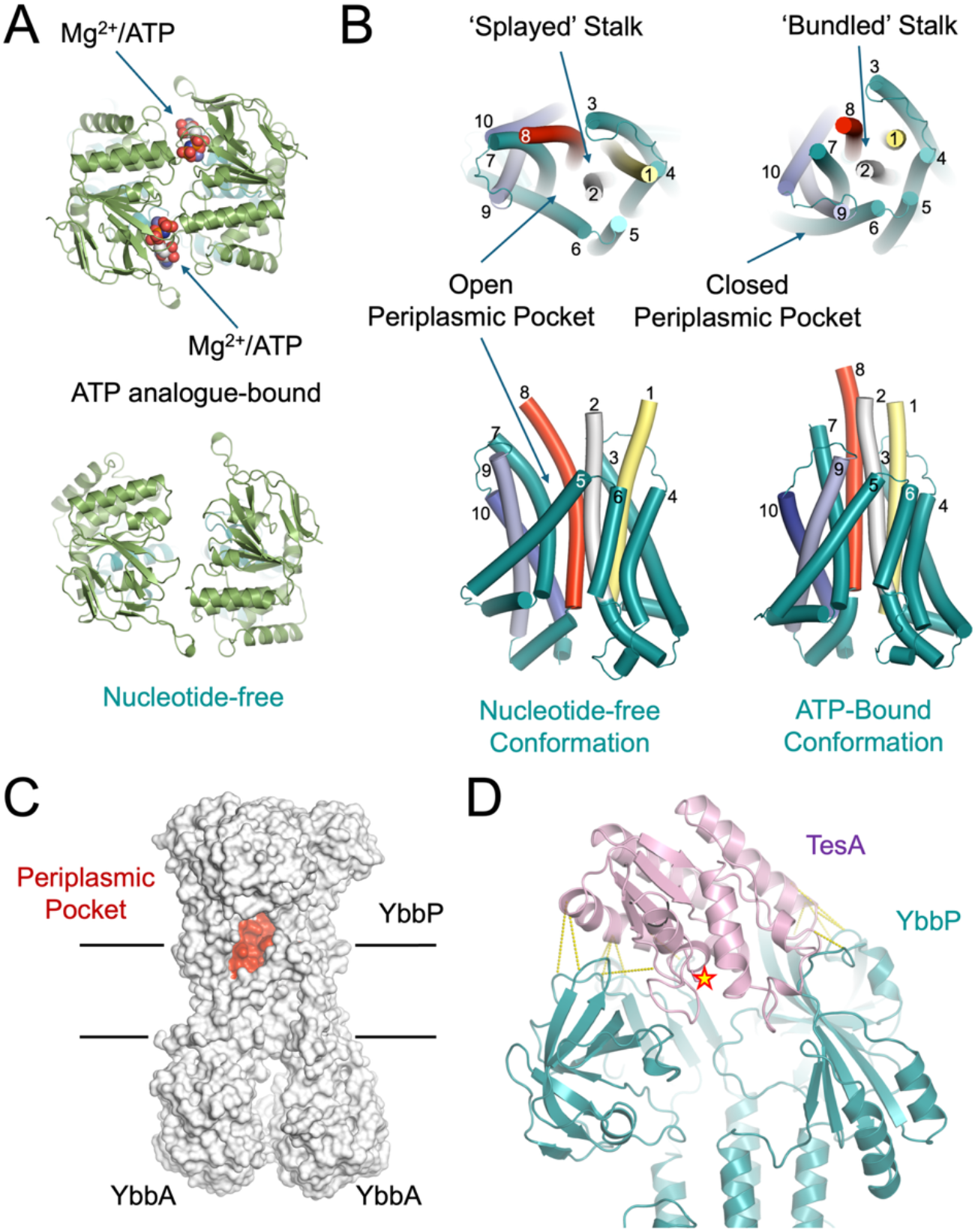
Evidence for a mechanotransmission mechanism in YbbAP-TesA. (a) Comparison of the YbbAP ABC domains in ATP-bound and nucleotide free structures. (b) Conformational changes in the YbbAP-TesA transmembrane domain. Transmembrane segments are numbered by their appearance in the amino acid sequence and both the periplasmic domains and ABC domains are omitted. To aid orientation, colours are used to highlight transmembrane helices 1, 2, 8, 9 and 10 which undergo large conformational changes. (c) Location of the Periplasmic-side pocket found in nucleotide-free YbbAP. (d) Coevolving residue pairs at the interface between YbbAP and TesA. Co-evolving residue pairs are shown with yellow dashes and the TesA active site is denoted with a star. A list of the top co-evolving residues is given in **Table S1**.

We identified three key differences between the structures of ATP-bound and nucleotide-free forms. Firstly, the YbbAP ABC domains are parted in the nucleotide-free structure and locked together in the ATP-bound forms (**Fig. 3a**). Secondly, the four helices on the periplasmic side of the membrane form a tight bundle in the ATP-bound conformation but are splayed apart in the nucleotide-free form causing large movements of the periplasmic domains in YbbP (**Fig. 3b**). Thirdly, we detected a large conformational change on the periplasmic side of the cytoplasmic membrane due to the repositioning of transmembrane helices 6 and 9 leading to the formation of a periplasmic-side pocket (**Fig. 3c**).

The scale of conformational change throughout the YbbAP complex is exceptionally large. For example, the distance between YbbA monomers is approximately 8.1 Å for both ATP binding sites (measured between Ser50 and Ser148), while the same distance measured in the absence of nucleotide is 17.2 Å or 23.6 Å (**Fig. 3a**). Similarly, the opening of the helical bundle on the periplasmic side of YbbAP involves large displacements that reversibly reveal and conceal the periplasmic-side pocket. In the nucleotide-free state, side-by-side packing between transmembrane helices 6 and 9 place Leu403 and Leu754 in close contact (6.6 Å between Cα positions) however, these residues are separated by 19.2 Å in the ATP-bound conformation due to the parting of these helices (**Fig. 3b**). The cavity formed inside the periplasmic pocket is hydrophobic in character and likely to be accessible to lipids from the outer face of the cytoplasmic membrane bilayer.

Consistent with other Type VII ABC systems, we find no evidence for the formation of a central pore or a pair of alternating access points that would allow passage of substrates across the membrane via YbbAP. The structural data strongly support the operation of a mechanotransmission mechanism in YbbAP-TesA in which ATP binding and hydrolysis drive periplasmic side conformational changes (especially opening and closing of the periplasmic pocket) rather than transport substrates across the inner membrane.

### The TesA active site is oriented to receive substrates from YbbAP

We next considered the orientation of TesA in relation to the YbbAP ABC transporter component. TesA is a relatively small protein in comparison to YbbAP and is located on top of the two periplasmic domains. The entrance to the TesA active site faces downwards towards the centre of YbbAP forming a substantial cavity that links the interior of TesA with that of the YbbAP periplasmic domains (**Fig. 3d**). Conserved TesA residues Asn138 and Tyr143 seem particularly important as they each form multiple contacts with residues in the YbbP periplasmic domain. Mapping the co-evolving residue pairs to the YbbAP-TesA cryoEM structure also confirms several important protein-protein contacts between TesA and both periplasmic domains of YbbP. These include electrostatic interactions between Arg141 of TesA and Asp148 of YbbP, and similarly, between Arg142 of TesA and Asp145 of YbbP. Other co-evolving residues highlight a mixture of polar and hydrophobic contacts that collectively serve to lock the hydrolase in place (**Fig. 3d** and **Table S1**). The TesA active site seems optimally positioned to receive substrates from YbbAP suggesting the transporter has a role in extracting compounds from the membrane for delivery to TesA.

### YbbAP-TesA has esterase activity towards p-nitrophenyl butyrate and nitrocefin, and thioesterase activity against Acetyl CoA

*E. coli* TesA is characterised as a multifunctional esterase with hydrolytic activity against several different fatty acyl substrates and lysophospholipids as well as thioesterase activity towards acetyl-CoA (37, 38, 43–45). *E. coli* TesA has also been suggested to have proteolytic activity (46), although this has not been detected in its *P. aeruginosa* homologue (38). To test whether the YbbAP-TesA complex retains the enzymatic activity of isolated TesA, we characterised YbbAP-TesA’s esterase activity after purification in detergent. We found that YbbAP-TesA has robust hydrolytic activity against p-nitrophenyl acetate, p-nitrophenyl butyrate and p-nitrophenyl octanoate (**Fig. 4a,b**). We also confirmed that YbbAP-TesA has thioesterase activity towards Acetyl CoA using Ellman’s reagent (47) to detect the release of free CoA-SH (**Fig. 4c**). These data confirm that YbbAP-TesA retains the same esterase (and thioesterase) activity as isolated TesA.

**Figure 4:**
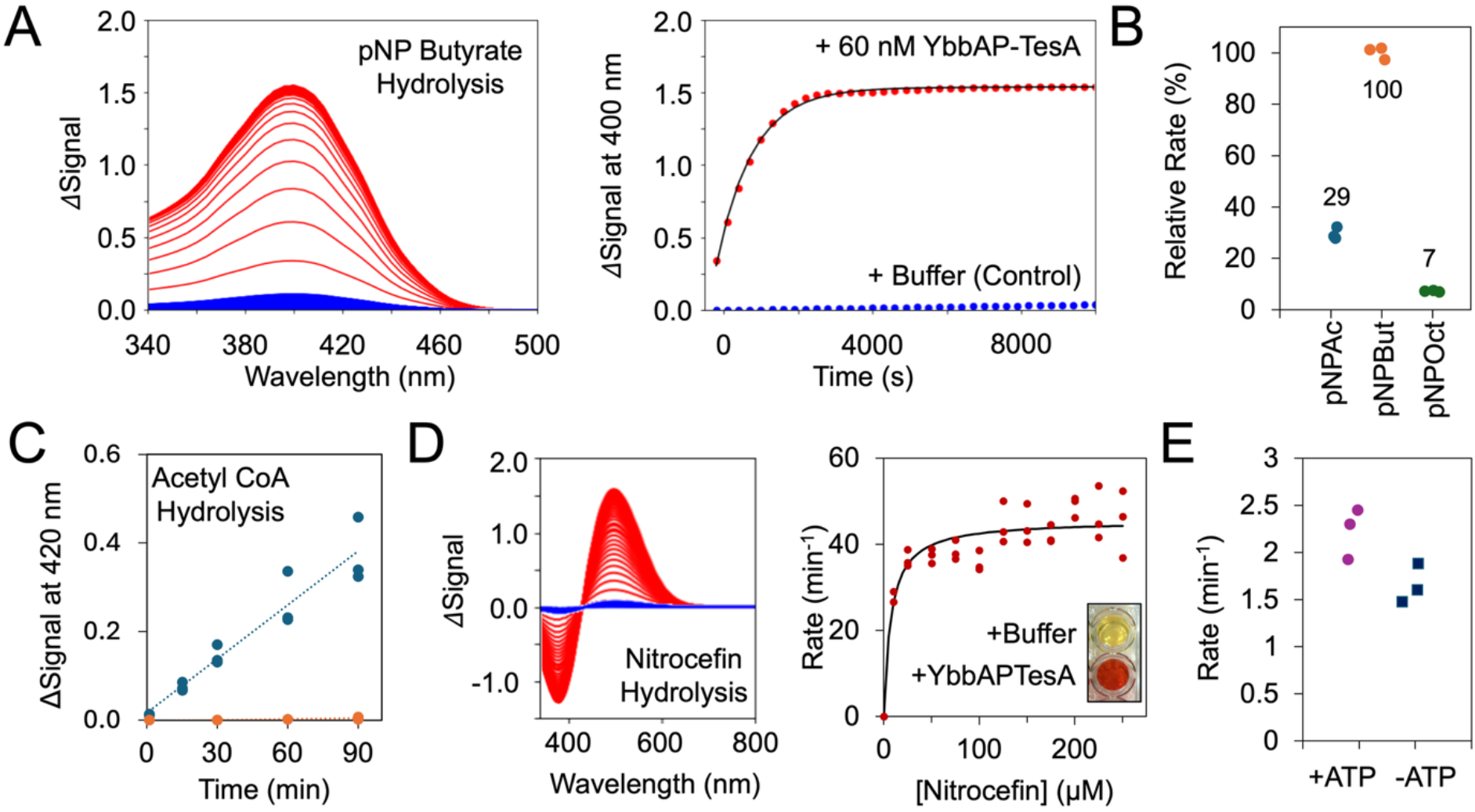
Esterase and thioesterase activity of YbbAP-TesA. (a) Hydrolysis of p-Nitrophenyl Butyrate by YbbAP-TesA produces a coloured product (p-Nitrophenol) that can be followed kinetically. The changes in absorption spectra are shown on the left and a plot of a single wavelength (400 nm) is shown verse time on the right. *Red* lines correspond to the enzyme-catalysed reaction and *blue* corresponds to buffer control. (b) Relative rates of hydrolysis for p-Nitrophenyl Acetate, p-Nitrophenyl Butyrate and p-Nitrophenyl Octanoate at 2.5 mM substrate concentration. Rates are normalised to the mean rate of p-Nitrophenyl Butyrate and given as a percentage. (c) YbbAP-TesA thioesterase activity monitored using Acetyl CoA as a substrate with detection of the CoA-SH product using DTNB. The signal at 420 nm is due to the production of thionitrobenzoate as DTNB reacts with the free thiol of CoA-SH. *Blue* dots indicate the enzyme catalysed reaction and orange dots represent the control. (d) Nitrocefin hydrolysis by YbbAP-TesA. A representative experiment showing changes in the absorbance spectra is shown on the left (250 μM nitrocefin) and a Michaelis Menten plot is shown on the right. A photo comparing the colour of two wells at the end of a 96-well plate experiment containing Nitrocefin solution with and without YbbAP-TesA is shown inset. (e) Rates of ester hydrolysis by YbbAP-TesA with and without 1 mM ATP. Experiments were performed at 21°C in PBS buffer with detergent, comparisons with and without ATP used 2.5 mM p-Nitrophenyl butyrate as the ester substrate.

In addition to these known substrates of TesA, we also found that YbbAP-TesA (and isolated TesA) can hydrolyse nitrocefin, a cephalosporin-like molecule commonly used to characterise beta lactamase enzymes (48) (**Fig. 4d**). While this finding initially led us to consider a role in low-level antibiotic hydrolysis, we found no evidence for enhanced resistance to beta lactam antibiotics when YbbAP-TesA was overexpressed, nor any loss of resistance when analysing YbbP or TesA knockout strains (**Table S3**). We therefore do not perceive a function for YbbAP-TesA in intrinsic resistance to beta lactams. Nonetheless, the colour change associated with nitrocefin hydrolysis makes nitrocefin a convenient substrate to further characterise YbbAP-TesA esterase activity. We measured a dissociation constant (*K*_*D*_) <10 nM for the hydrolysis of nitrocefin by YbbAP-TesA (**Fig. 4d**). We also separately confirmed that the *P. aeruginosa* TesA can hydrolyse nitrocefin (**Fig. S4**) showing that nitrocefin hydrolysis is likely to be a general property of the TesA family and not a peculiarity of *E. coli* YbbAP-TesA.

Frustratingly, we have not yet identified any differences between the substrate profiles of isolated TesA and the YbbAP-TesA complex. We also find little evidence for stimulation of YbbAP-TesA activity upon addition of ATP (**Fig. 4e**). This is most likely because the substrates used here are accessible to TesA without the need for extraction from the membrane by YbbAP. We therefore conclude that YbbAP-TesA retains the broad-spectrum esterase and thioesterase activity of TesA.

### *In vivo* activity of YbbAP-TesA towards p-nitrophenyl butyrate and identification of catalytic residues

To further characterise YbbAP-TesA esterase activity, we monitored p-nitrophenyl butyrate hydrolysis by live *E. coli* cells. Strains lacking *tesA* retain low levels of esterase activity above the level of spontaneous hydrolysis seen in controls absent of *E. coli*. However, the same strains expressing either TesA (or YbbAP-TesA) from a plasmid show robust hydrolytic activity towards p-nitrophenyl butyrate (**Fig. 5a**). As expected, expression of a TesA variant lacking the nucleophilic serine residue (Ser36Ala) does not restore hydrolase activity to wild type levels and neither does mock expression using an empty vector as a negative control.

**Figure 5:**
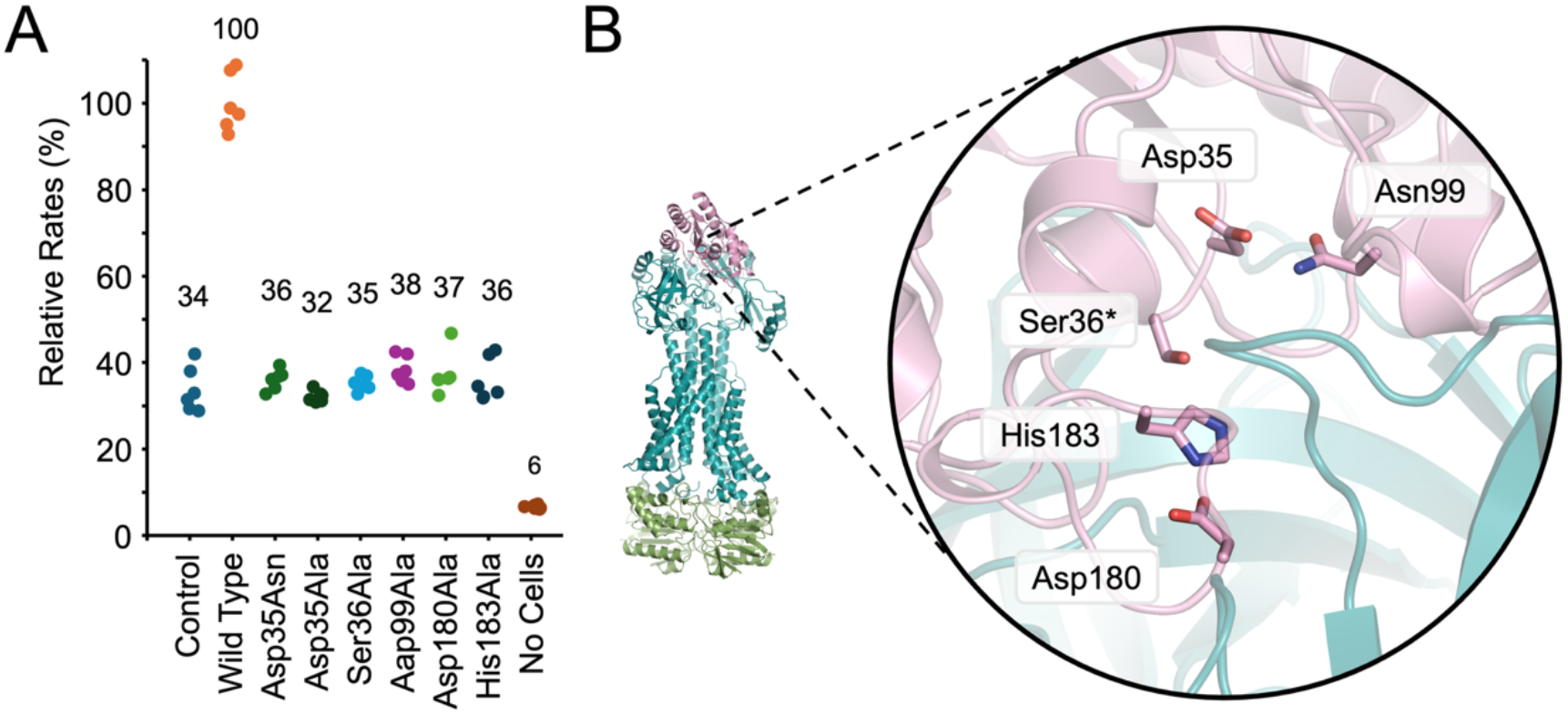
Mutational analysis of the TesA active site and *in vivo* esterase activity. (a) Relative *in vivo* esterase activity towards p-nitrophenyl butyrate. Control indicates the rate of p-nitrophenyl butyrate hydrolysis by *E. coli ΔtesA* cells carrying an empty pET vector. Other lanes indicate rates for *E. coli ΔtesA* cells carrying a pET vector expressing either ‘Wild Type’ TesA or the indicated TesA active site variant. Relative rates were measured six times (two biological repeats with three technical repeats each) and normalised to the wild type. (b) Location of proposed catalytic residues within the TesA active site.

We therefore made mutations in conserved TesA residues that are likely to be involved in catalysis. Mutations designed to impair catalysis were chosen based on previous TesA literature, structures of isolated TesA (38, 49), and inspection of our cryoEM structure (**Fig. 5b**). We also made use of residue conservation data from a large multiple sequence alignment using TesA homologues (**Fig. S5**). His183 and Asp180 form a highly conserved dyad that has been proposed to activate Ser36 by deprotonating its hydroxyl group. Consistently, we found that His183Ala and Asp180Ala are both catalytically impaired to the level of the negative control *in vivo* (**Fig. 5a,b**). We also mutated Asp35 which is part of the conserved GDSL motif from which this family of hydrolases has been named (40). Asp35 is located about 5 Å from the catalytic serine and forms a hydrogen bond to another highly conserved residue, Asn99. The roles of Asp35 and Asn99 are unclear but could conceivably involve substrate binding or acid-base catalysis. We found that Asp35Ala, Asp35Asn and Asn99Ala mutations all inactivate TesA activity suggesting these residues are required for function in addition to Ser36, His183 and Asp180 (**Fig. 5a**,**b**). These experiments confirm the essential catalytic residues within TesA.

### Molecular dynamics simulations suggest lysophospholipids access the periplasmic pocket in YbbAP

To investigate the hypothesis that YbbAP-TesA extracts substrates from the bacterial inner membrane, we conducted molecular dynamics simulations. The cryoEM structures of apo and ATP-bound YbbAP were first embedded in model membranes composed of *E. coli* lipids, converted to coarse-grained models, and simulated three times over 15 μs (**Fig. 6a,b**). The simulation setup is shown in **Fig. 6a**. Each membrane was supplemented with lysophospholipids (LysoPG and LysoPE) as candidate substrates for YbbAP and their locations were monitored over the course of the simulation (**Fig. 6b** and **Fig. S6**). In support of the proposed role for YbbAP in extracting lipids for presentation to TesA, the simulations revealed instances where lysophospholipids entered and occupied the periplasmic-side pocket of apo YbbAP (**Fig. 6b**). Residency plots for lipids in the upper leaflet of the membrane confirm accumulation of LysoPG within the periplasmic-side pocket (**Fig. 6b** *left*). As expected, lysophospholipid binding to the periplasmic side pocket was not observed for simulations that started from the ATP-bound state where the pocket is closed (**Fig. 6b** *right*).

**Figure 6:**
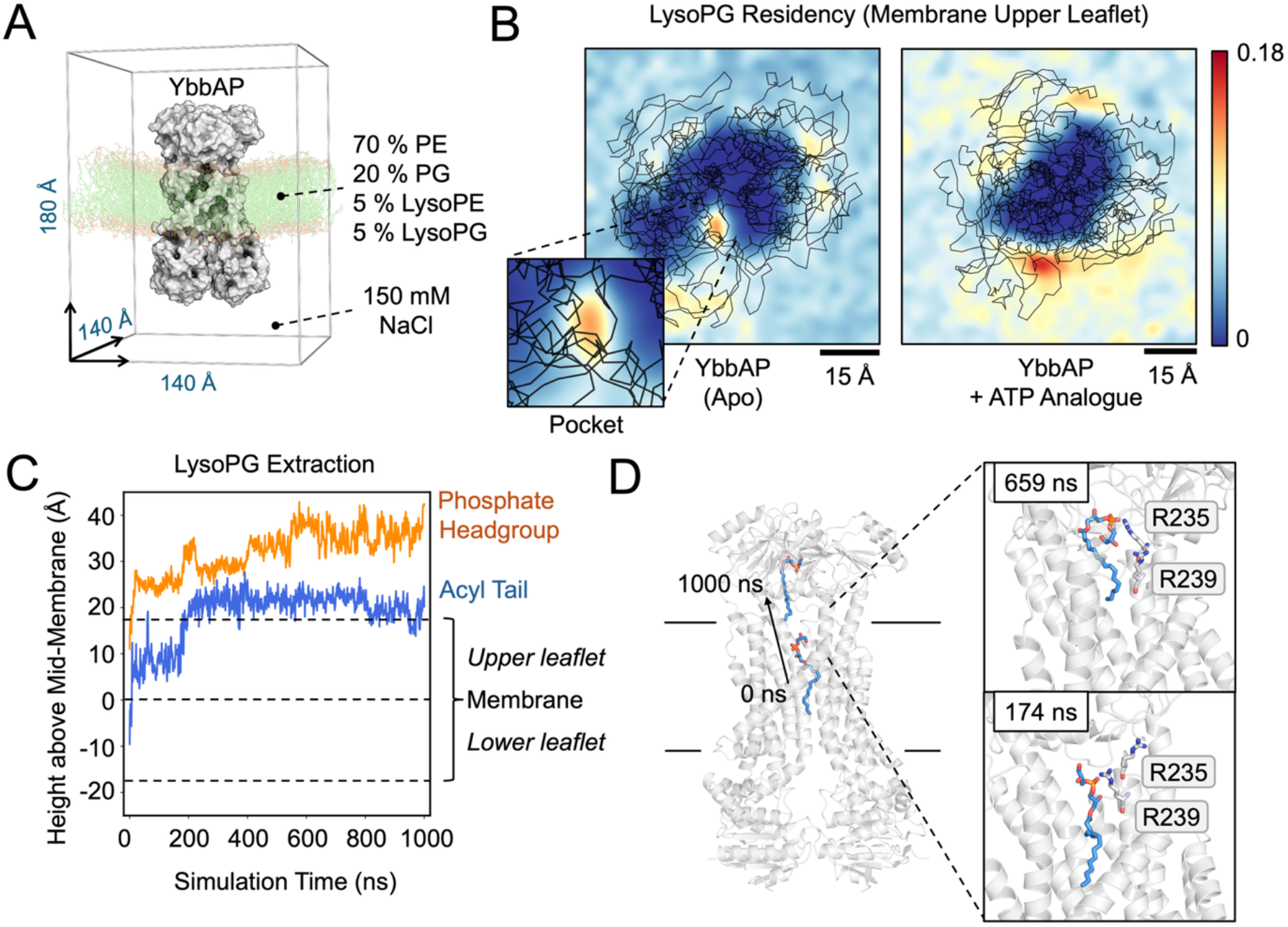
Molecular dynamics simulations of YbbAP suggest the periplasmic-side pocket is a lipid binding site. (a) Simulation setup. (b) 2D histogram (bins: 30×30) of LysoPG phosphate head group residency in the membrane upper leaflet over 15 μs coarse-grain simulation repeats of YbbAP in its apo and nucleotide-bound states. Scale is normalised to the global maximum. A close-up view of the area encompassing the periplasmic pocket is shown inset. Protein backbone coordinates are shown in black. Residency is indicated by the frequency density of LysoPG headgroups occupying each 2.5 Å by 2.5 Å area. The highest observed frequency density across both datasets was 0.18 (n = 6, 3 per structure), from a trajectory of frames comprising of every 10^th^ 1 ns simulation time point. Densities of LysoPE and other lipids are given in **Supplemental Figure S6**. (c) Vertical movement of a single LysoPG molecule that undergoes extraction from the bilayer over 1000 ns of atomistic simulation, starting from the end point of the coarse-grain simulation where LysoPG occupies the periplasmic pocket. Repeats in **Supplemental Figure S8**. (d) Key frames from the start and end of the atomistic simulation showing LysoPG exiting the bilayer. Residues that appear to assist LysoPG extraction during the atomistic simulation are indicated.

To further explore the behaviour of lyosphophospholipids that entered the periplasmic-side pocket, we performed additional molecular dynamics simulations using fully atomistic models (**Fig. 6c,d** and **Fig. S7**). Using the final frame of our coarse-grained simulation as a start point, we followed the movement of a pocket-bound lysophospholipid (LysoPG) over 1000 ns. In two of the three simulations, LysoPG was partially extracted from the bilayer by YbbAP with both the phosphate headgroup and acyl tail rising out of the membrane and into the periplasmic domain of YbbP (**Fig. 6c** and **Fig. S8**). Further inspection of the atomistic simulations (including protein-lipid contact analysis – **Fig. S8b**) revealed that two arginine residues (Arg235 and Arg239) are likely to be important in extracting lysophospholipids as they form complementary electrostatic interactions with the phosphate of the lysophospholipid head group (**Fig. 6d**).

Association of lysophospholipids with the YbbAP periplasmic-side pocket and knowledge of the different conformational states observed by cryoEM suggests a mechanism for lipid extraction by YbbAP-TesA. Specifically, we propose that substrates first bind to the periplasmic side pocket, in ADP-bound/nucleotide free conformation of YbbAP. Closure of the lipid-occupied periplasmic pocket is then driven by ATP binding. This conformational rearrangement extracts the lipid from the membrane through mechanotransmission and forces the contents of the pocket upwards towards and then into the TesA active site. Hydrolysis of ATP likely causes the system to reset, allowing for further rounds of substrate binding and delivery to the TesA active site.

## Discussion

Type VII ABC transporters are bacterial membrane protein complexes that typically pair an ABC transporter with a periplasmic partner protein to establish their biological function. Here we used computational methods to identify TesA as the partner of YbbAP and purified the YbbAP-TesA complex from *E. coli* (**Fig. 1**). A cryoEM structure of YbbAP-TesA shows that TesA is located at the interface between the two periplasmic domains of YbbP with its active site oriented to receive substrates from the ABC transporter (**Fig. 2**). Structures of YbbAP with and without a bound ATP analogue also capture dramatic transmembrane conformational changes that are the signature of a mechanotransmission mechanism (**Fig. 3**). *In vitro* activity assays show that YbbAP-TesA has hydrolytic activity against several fatty acyl-esters, acetyl-CoA and nitrocefin (**Fig. 4**). Mutagenesis of the active site also confirms a constellation of residues that are essential for *in vivo* esterase activity (**Fig. 5**) and simulations of YbbAP-TesA in model membranes predict that lipids may directly interact with a periplasmic pocket close to the YbbAP-TesA active site (**Fig. 6**). These results are consistent with a role for YbbAP in extracting lipids and other hydrophobic compounds from the bacterial inner membrane for hydrolysis by TesA.

YbbAP-TesA is the fourth Type VII ABC transporter system to be characterised in *E. coli* (**Fig. 7**). The three other Type VII ABC systems each have important roles in the bacterial cell envelope including cell division (FtsEX-EnvC), antibiotic resistance and toxin secretion (MacAB-TolC), and lipoprotein transport (LolCDE-LolA). While it has been clear for some time that YbbAP is structurally and functionally distinct from these other Type VII ABC systems (2), the lack of a distinctive phenotype upon genetic deletion has delayed its characterisation. Understanding that YbbAP is part of a larger complex with TesA has now led us to propose a role for YbbAP-TesA in extracting and hydrolysing hydrophobic compounds from the bacterial inner membrane. While the natural substrates of TesA are not completely clear, the enzyme unambiguously has native esterase and thioesterase activity and association with YbbAP likely signals a connection with substrates in the lipid bilayer. Observation of large periplasmic-side conformational changes driven by cytoplasmic side ATP binding and hydrolysis in the YbbAP structures are consistent with a mechanotransmission mechanism and could form a structural basis for extraction of lipid-like substrates (or other hydrophobic molecules) from the membrane for presentation to the TesA active site.

**Figure 7:**
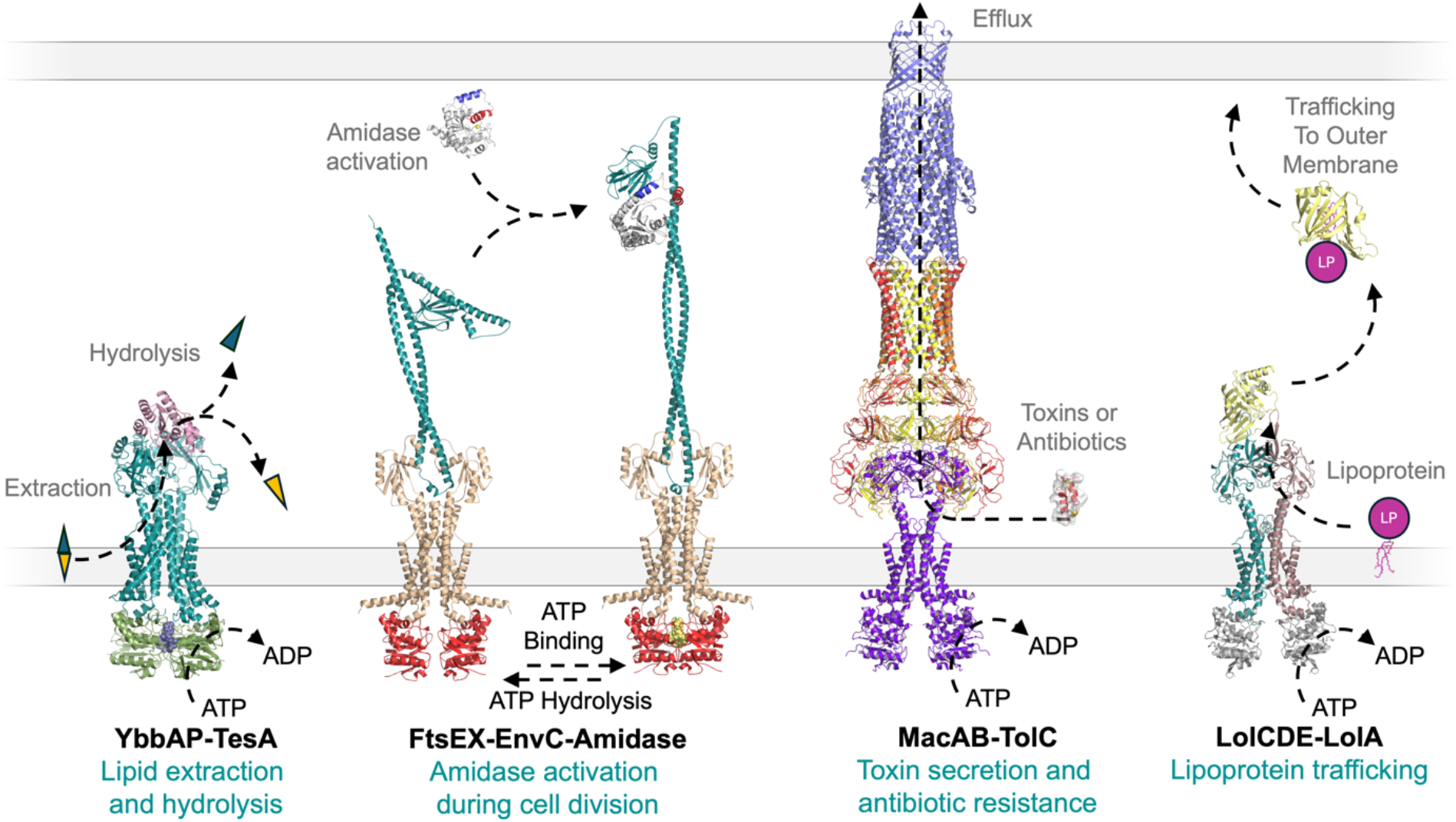
Biological functions of the four Type VII ABC transporters in *E. coli*. From left-to-right: YbbAP-TesA forms a novel type VII ABC transporter involved in extraction of hydrophobic compounds from the inner membrane and enzymatic hydrolysis in the periplasm; FtsEX-EnvC uses transmembrane conformational changes to regulate the activity of peptidoglycan amidases in the periplasm; MacAB-TolC drive efflux of antibiotics and small toxins across the outer membrane; and LolCDE extracts lipoproteins from the inner membrane and delivers them to the outer membrane via the LolA shuttle.

The two most likely natural substrates for YbbAP-TesA are lysophospholipids and Acyl-CoA as these are the only natural biomolecules that have been shown to be substrates of the TesA hydrolase (37, 38, 43–45). Lysophospholipids are thought to be toxic due to their destabilising effects on membrane integrity (50). Lysophospholipids are produced endogenously during lipoprotein maturation by the activity of lipoprotein N-acyltransferase (Lnt) (51–53) and can also be generated by secreted enzymes of the immune system (54, 55) or injected effectors of bacterial competitors (50, 56). It is therefore plausible that YbbAP-TesA has a role in detoxifying (or recycling) lysophospholipids that might otherwise accumulate in the bacterial cell membrane. Alternatively, hydrolysis of Acetyl CoA (or membrane partitioning Acyl-CoA) might provide access to useful biochemical materials such as Acetate, Coenzyme A and fatty acids.

In summary, we have isolated and characterised the *E. coli* YbbAP-TesA complex; a new Type VII ABC transporter system that appears to extract molecules from the bacterial inner membrane and presents them to an attached periplasmic hydrolase for hydrolysis. Our study shows the remarkable diversity of functions among bacterial ABC transporters and demonstrates how computational methods and experimental approaches can be effectively combined to identify previously uncharacterised Type VII ABC transporter systems.

## Methods

Complete methods are provided in the **Supplementary Appendix**. Briefly, YbbAP-TesA complexes were purified from *E. coli* C43(DE3) after co-expression from pETDuet1-based plasmids using Ni-affinity chromatography with a His-tag on YbbA. Samples used for structure determination were further purified using size exclusion chromatography. Membrane protein complexes were stabilised in 0.01 % LMNG detergent and ATP-analogues (AMP-PnP; Adenosine 5′-(β,γ-imido)triphosphate) added as required. For CryoEM, protein samples were frozen on Quantifoil 300 mesh grids using an FEI Mark IV Vitrobot and data was collected using a Titan Krios electron microscope. Structures of YbbAP-TesA bound to an ATP-analogue and YbbAP (with and without an ATP-analogue) were determined using software from Relion (57) and Phenix (58). Prospective partner proteins for YbbAP were identified computationally using a brute-force co-folding approach implemented with Colabfold (35, 36) and inter-protein coevolution was assessed using CCMPred (42). *In vivo* activity assays used a TesA knockout strain from the Keio collection (59) complemented with either TesA, YbbAP-TesA or various TesA single amino acid variants expressed from a plasmid. Cells were grown in LB with 50 μg/mL ampicillin and induced with 1mM IPTG, before being resuspended at OD 1 in phosphate buffered saline (PBS) and added to an equal volume of 5mM p-nitrophenol butyrate in PBS. Rates were determined using absorbance at 420 nm. Enzyme assays were performed in 96-well plates at 21°C in PBS buffer supplemented with 0.01% LMNG. Experiments using p-Nitrophenol butyrate, p-Nitrophenyl octanoate, and p-Nitrophenyl acetate were followed kinetically at 400 nm and nitrocefin at 496 nm, while AcetylCoA hydrolysis was measured at intervals after stopping the thioesterase reaction using 8 M Urea and 2 mM DTNB. Molecular dynamics used GROMACS (60) for both coarse-grained and atomistic simulations with MEMEMBED (61) and INSANE (62) used for simulation setup. The CHARMM forcefield was used for atomistic simulations (63). Lipid density was analysed with PLUMED and data further analysed using MDAnalysis (64, 65). Structures and simulations were visualised with Pymol (66).

## Data, Materials and Software Availability

CryoEM structures have been deposited in the Protein Data Bank under accession codes 9GE6, 9GE7 and 9GE8. The corresponding cryoEM maps are deposited in the Electron Microscopy Data Bank with accession numbers EMD-51291, EMD-51292 and EMD-51293.

## Acknowledgements

We thank colleagues at the University of Leicester for support with CryoEM sample preparation and data processing, the School of Life Sciences media kitchen for supply of bacterial growth media and the Tech Team for laboratory support. We thank Daisy Robinson for undergraduate project work exploring phenotypes of YbbP knockouts and Prof. Freya Harrison for providing *P. aeruginosa* genomic DNA. We acknowledge use of computational resources on Archer2 (HECBioSim, EPSRC EP/R029407/1), Sulis (HPC Midlands+, EP/T022108/1) and the University of Warwick Scientific Computing Research Technology Platform. The work was funded by MRC-DTP (MR/N014294/1), BBSRC-MIBTP (BB/T00746X/1), Wellcome (208361/Z/17/Z), MRC, BBSRC, EPSRC, NIH, JPIAMR, the University of Warwick and the Howard Dalton Centre.

