## Supplemental Information for "Structure of YbbAP-TesA: a Type VII ABC transporter–lipid hydrolase complex"

**Schedule of Supplementary Data**

**Supplementary Methods**

**Supplemental Tables**

Table S1: Co-evolving residues predicted between YbbA, YbbP and TesA
Table S2: CryoEM data and model refinement statistics
Table S3: Antibiotic susceptibility testing for TesA and YbbP knockout strains

**Supplemental Figures**

Figure S1: Image processing workflow for the AMP-PNP bound YbbAP cryo-EM reconstruction.
Figure S2: Image processing workflow for the apo YbbAP cryo-EM reconstruction.
Figure S3: Image processing workflow for the AMP-PNP bound YbbAP-TesA complex cryo-EM reconstruction.
Figure S4: Nitrocefin and p-nitrophenyl butyrate hydrolysis assays for *P. aeruginosa* TesA
Figure S5: Amino acid sequence conservation in TesA
Figure S6: RMSF and RMSD calculations for atomistic simulations of YbbAP.
Figure S7: Lipid residency distributions during coarse-grain simulations of YbbAP.
Figure S8: LysoPG trajectory in atomistic simulation of YbbAP.

**Movie Legends**

Movie 1: Evidence for a mechanotransmission mechanism in YbbAP-TesA.

**Supplemental References**

### Methods

**Cloning, expression and purification of YbbAP and YbbAP-TesA**

For expression of YbbAP-TesA, the YbbAP gene pair was cloned into the first multiple cloning site of pETDuet-1 in frame with the N-terminal His-tag and TesA was cloned into the second multiple cloning site without a tag. The resulting vector (pET/YbbAP/TesA) was transformed into *E. coli* C43(DE3) for expression. Small scale cultures were first grown in Lysogeny Broth (LB) and then transferred to 2YT for expression at scale (typically 6 L of media divided among six 2.5 L flasks). Cells were grown at 30°C in 2YT media supplemented with 50 µg/mL ampicillin, induced by addition of 1 mM IPTG and allowed to express proteins overnight at 18 °C. Cells were harvested by centrifugation in 1 L flasks (10,000 ×g, 10 min at 10 °C) and the pellets frozen in 50 mL falcon tubes at -80 °C until required. Cells were thawed at room temperature and resuspended in a buffer composed of 30 mM Imidazole, 300 mM NaCl, 50 mM HEPES pH 7.2 ('Wash buffer'). Resuspended cells were then supplemented with lysozyme and DNase I and broken using multiple passes through an Avestin EmulsiFlex-C3 at 15,000 psi. Unbroken cells and debris were removed from the resultant cell lysate using a low-speed centrifugation step (10,000 ×g, 10 min, 10 °C). The supernatant was then centrifuged at 100,000 ×g for 1.5 h at 6 °C using a Sorval ultracentrifuge to pellet membranes. The supernatant was discarded and membranes resuspended in fresh buffer. Membranes were solubilised by the addition of detergent (0.5 % LMNG). After 2h, the sample was ultracentrifuged again to remove insoluble material (100,000 ×g, 45 min, 6 °C). Detergent-solubilised proteins were incubated with Ni-IMAC resin overnight and transferred to an empty PD-10 column. The protein-bound resin was washed with Buffer A supplemented with 0.005 % LMNG and eluted from the resin using a buffer composed of 250 mM Imidazole, 300 mM NaCl, 50 mM HEPES pH 7.2 (Elution Buffer) supplemented with 0.005 % LMNG. Proteins were concentrated using centrifugal filter devices with a nominal 100 kDa molecular weight cutoff. A pre-packed PD10 Gel filtration column was used to exchange buffers for enzymatic studies (PBS supplemented with 0.01 % LMNG). Samples destined for enzymatic study were supplemented with 10 % glycerol and flash frozen in aliquots using liquid nitrogen and stored at -70 °C for use as required.

For cryoEM, the YbbAP-TesA(Ser36Ala) protein was produced similarly to the wild type using the pETDuet/YbbAP/TesA(Ser36Ala) expression vector, however, a size exclusion chromatography step was employed after elution from the Ni-IMAC resin in place of the buffer exchange. Specifically, concentrated YbbAP-TesA(Ser36Ala) was injected onto a Superdex 200 increase (10/300) column pre-equilibrated with a buffer composed of 20 mM HEPES, 150 mM NaCl, 0.005% LMNG pH 7.2 and fractions corresponding to the main peak were collected and pooled. CryoEM samples were made fresh and never frozen prior to grid preparation. The pETDuet/YbbAP/TesA(Ser36Ala) expression vector was constructed similarly to the pET/YbbAP/TesA plasmid using the native YbbAP operon in the first Multiple Cloning Site (MCS) and a Genscript-synthesised TesA(Ser36Ala) variant in the second.

The YbbAP protein was produced similarly to YbbAP-TesA using plasmid pET/YbbAP. The pET/YbbAP construct was obtained by cloning the YbbAP gene pair into pETDuet via the BamHI and XhoI restriction sites. The YbbAP genes therefore span both cloning sites and encode a His-tagged YbbA proteins and untagged YbbP.

**Cloning, expression and purification of soluble TesA**

Soluble His-tagged *E. coli* TesA (with a C-terminal His tag) was produced using the pET/TesA expression vector in *E. coli* C43(DE3) cells. pET/TesA was constructed by cloning the DNA encoding residues 27-208 into pET21a via NdeI/XhoI sites. Cells were grown at 30 °C in 2YT, and induced with 1 mM IPTG for 18h at 30 °C. After expression, cells were pelleted by centrifugation (10,000 ×g, 10 min at 10 °C), the supernatant was removed and pelleted cells and stored at -80 °C in 50 mL tubes. Cells were thawed, resuspended in Wash buffer (20 mM Imidazole, 300 mM NaCl, 50 mM HEPES pH 7.2) and then supplemented with DNase I and lysozyme before being broken by sonication. Cellular debris and unbroken cells were removed by centrifugation (30,000 ×g, 20 min at 10 °C). The cleared lysate was incubated overnight with Biorad Ni-IMAC resin and then passed through an empty PD10 column, washed with Wash buffer, and then eluted using 2.5 mL of Elution buffer (250 mM Imidazole, 300 mM NaCl, 50 mM HEPES pH 7.5). Proteins were then buffer exchanged into PBS using a PD10 Gel filtration column. Proteins used for enzyme assays were supplemented with 10 % glycerol, divided into aliquots and frozen at -80°C for later use. The *P. aeruginosa* TesA was cloned and produced similarly to *E. coli* TesA using *P. aeruginosa* PA14 genomic DNA as a PCR template.

**Enzyme assays**

AcetylCoA was solubilised in PBS buffer at 13 mM and frozen in aliquots for use as required. Enzyme assays used AcetylCoA at 1 mM in PBS buffer supplemented with 0.01% (w/v) LMNG detergent. Reactions were started by the addition of enzyme (or 0.01 % LMNG in PBS buffer as a negative control) in equal volume at 0.6 μM. For each time point, a 100 μL volume was removed from the reaction mixture and added to an equal volume of 'Stopping buffer' composed of 8 M Urea and 2 mM DTNB (Ellman's reagent) in PBS. The absorbance spectrum was then measured in a 96-well plate reader. An additional control reaction containing the enzyme, but no Acetyl CoA substrate was used to show that DTNB's reaction with protein thiols is insignificant compared to the reaction with substrate.

P-nitrophenyl acetate, p-nitrophenyl butyrate and p-nitrophenyl octanoate were obtained from Sigma. Solutions involving p-nitrophenyl derivatives were freshly prepared immediately before use. Hydrolysis of p-nitrophenyl derivatives was conducted in PBS buffer supplemented with 0.01 % LMNG. Enzymes were purified as above with *E. coli* TesA, *P. aeruginosa* TesA or the YbbAP-TesA complex purified in PBS or PBS with 0.01 % LMNG. Where necessary, the buffer was supplemented with 1mM $MgCl_2$ and ATP was added at 1 mM. Reactions were started by addition of enzyme (or enzyme buffer) and followed kinetically, in parallel, using a plate reader at 21 °C. Controls were performed identically with the addition of buffer in place of enzyme. The starting point absorbance measurement from the control was used for background subtraction.

Nitrocefin was prepared as a 13 mM stock in PBS and frozen in aliquots before use. Nitrocefin hydrolysis was followed by its change in colour over time using a plate reader. Nitrocefin hydrolysis assays were performed in PBS buffer supplemented with 0.01 % LMNG at 21 °C and started by the addition of enzyme. Where possible, rates were derived by fitting the kinetic data to a single exponential equation, or else using the initial rates and the product extinction coefficient.

##### Computational prediction of partner proteins using *in silico* co-folding

A list of candidate periplasmic protein sequences was identified using Uniprot (57) and the sequence of YbbP retrieved from KEGG (41). Candidate proteins were then co-folded pairwise with YbbP using Colabfold (36) before calculating the pDockQ (34) and Molprobity (39) clash scores for each predicted complex.

##### Coevolution analyses

Amino acid sequences for TesA, YbbA and YbbP were obtained from the *E. coli* K-12 genome via KEGG (41). Paired alignments of homologous sequences were then produced using

ColabFold (36). Co-evolution within the paired alignment was analysed using CCMPred which assigns a pairwise score to every position in the paired alignment (42). The mean and standard deviation for all *intermolecular* pairs was calculated and used to assign Z-scores for each individual amino acid pair. Histograms of the intermolecular coevolution Z-score values and 2D coevolution plots were constructed and visualised using the Matplotlib. Co-evolving residue pairs with high Z-scores were mapped to structures using Pymol (58) and residue conservation diagrams were created with WebLogo (59).

**CryoEM data collection, processing and model building**

Quantifoil 300 mesh gold grids (1.2/1.3 Quantifoil Micro Tools EMBH) were preprepared by glow discharge for 60 seconds at 35 mM using a Quorum GloQube instrument. Protein samples concentrated to approximately between 0.5 mg/mL and 0.9 mg/mL were then applied to the grids before blotting for 3 seconds and plunge freezing in liquid ethane using an FEI Vitribot Mark IV (blot force 10, 100% humidity). Vitrified grids were screened with an FEI Titan Krios G3 and suitable samples were selected for data collection. Data was captured using a K3 detector and BioQuantum energy filter with 20 eV slit width. The dose rate was 17 e $pixel^{-1}$ $s^{-1}$ with a total dose of 49 e $Å^{-2}$ and fractionated at 1 e $Å^{-2}$. Collections were prepared and automated using EPU2 software across a defocus range of -0.8 to -2.3 µm.

All data processing was completed using Relion 5 (60) but initiated using Relion 4.0 (61, 62). Micrographs were motion-corrected using MotionCor2 and CTF estimation was performed using CTFFIND4 (63). Micrographs were manually selected for good CTF fits before subjecting to particle picking used the Relion implementation of Topaz (64) which was trained using sub-1,000 manually-picked particles. Exact processing methods varied between datasets and are detailed in **Figures SI3-5.** Particles were initially extracted and down-sampled for rounds of initial 2D classifications, ab initio model generation and 3D classifications. For the initial 3D classification, 60 Å low-pass filtered initial models were used as references with subsequent 3D auto refined models used as reference for further 3D classifications, each with reduced low-pass filtering. For 2/3 datasets, particles were re-extraction at the full resolution of 0.835 Å $pixel^{-1}$ and subjected to rounds of CTF-refinement and Bayesian polishing before a final auto-refinement with Blush regularisation (65). For the smallest dataset of purified YbbAP-TesA complex, particle numbers were limited and so further 3D classifications were performed against all picked particles to select more good particles before CTF refinement and polishing. Sharpened maps were produced using Phenix.autosharpen (66, 67). Where required, the handedness of cryoEM maps was reversed using ChimeraX (68).

Initial 3D structural models were produced by Alphafold3 (69) with cycles of manual model rebuilding using Coot (70) and refinement using Phenix.Refine (71) to complete each model. Late-stage model building was informed by validation tools from Coot and Molprobity (39, 70). Coordinates and cryoEM maps for the three structures have been deposited in the Protein Data Bank (72) with accession codes 9GE6 (YbbAP), 9GE7 (YbbAP with bound ATP analogue) and 9GE8 (YbbAP-TesA with bound ATP analogue).

**Antibiotic susceptibility assays**

Minimum Inhibitory Concentrations were determined for *E. coli* BW25113 (parental strain), BW25113 *Δtesa*::*kan* (strain lacking TesA) and BW25113 *Δybbp*::*kan* (strain lacking YbbP) using penicillin G, cephalothin, and cephalexin. MICs were performed in 96-well plates using LB broth and 2-fold dilutions. Growth at 37 °C was recorded using the OD 600 after 18h. MICs were also determined for *E. coli* C43(DE3) carrying pET28/YbbAP/TesA, pET28/YbbAP/TesA(S36A) and pET28/YbbAP using similar methods (37 °C, 48h), except LB was supplemented with 25 µg/mL Kanamycin to maintain plasmid selection and 1 mM IPTG to induce expression of plasmid-encoded proteins. The pET28/YbbAPTesA and pET28/YbbAP constructs were generated by subcloning NcoI/XhoI digested DNA fragments from pET/YbbAP or pET/YbbAP/TesA amnd ligating into similarly digested pET28a.

***In vivo* activity assays**

*E. coli* strains lacking TesA were obtained from the Keio collection (73) and transformed with pET-based vectors designed to express full length TesA or TesA variant. The parental *E. coli* strain BW25113 and *E. coli ΔtesA*::*kan* were used as comparators and were transformed with the corresponding empty vector. Cells were grown in LB supplemented with 50 µg/mL ampicillin, induced with 1mM IPTG, before twice being pelleted (6000 ×g) and then resuspended at OD 1 in PBS buffer. A reaction mixture containing 5 mM p-nitrophenyl butyrate in PBS was then transferred into 96-well plates ready to receive live bacterial cells. The assay was started by addition of one third volume of resuspended bacterial cells and followed by spectrophotometric measurements at 15 min intervals. The experiment was repeated three times and then again using freshly transformed cells (ie two biological repeats, with three technical repeats each). Relative rates were estimated using absorbance at 420 nm vs time.

**Molecular dynamics simulations**

The membrane orientation for each cryoEM structure was predicted using MemEmbed (74), and the resulting models were centred in a cuboid box and coarse-grained to the Martini3 representation using Martinize2 (75). Coarse-grained lipid membranes consisting of 70% 1-palmitoyl-2-oleoyl-sn-glycerol-3-phosphotidylethanolamine (PE), 20% 1-palmitoyl-2-oleoyl-sn-glycerol-3-phosphotidylglycerol (PG), 5% 1-palmitoyl-sn-glycerol-3-phosphotidylglycerol (LysoPG), and 5% 1-palmitoyl-sn-glycerol-3-phosphatidylethanolamine (LysoPE) were assembled around each protein using *insane* (76), and each system was solvated with water and 0.15 M NaCl. Systems were energy minimised using the steepest descent algorithm, and a two-step equilibration which applied velocity-rescaling temperature coupling and a semi-isotropic Berendsen barostat (77, 78). Three production runs were then performed by applying a semi-isotropic c-rescale barostat at 1 bar and v-rescale temperature coupling at 310 K to each equilibrated system over a simulation length of 15 μs (79). All coarse-grained simulations were run using GROMACS 2022.3, electrostatic terms were described using reaction field electrostatics with a cut-off of 1.1 (80). Van der Waals interactions were calculated using a 1.1 nm cutoff. A timestep of 20 fs was used for all CGMD simulations.

The final frame of the coarse-grained simulation of Apo-YbbAP showing LysoPG occupation of the binding pocket was converted to atomistic representation using CG2AT2 (81). Three simulation systems were generated from this single frame, using the CHARMM36m forcefield and TIP3P water (82–84). Energy minimisation was carried out using the steepest descents algorithm. A 10 ns run was performed with position restraints applied to the protein, followed by a 1 μs production run using a 2 fs timestep and a semi-isotropic c-rescale barostat at 1 bar and v-rescale temperature coupling at 310 K. Hydrogen bonds were constrained using Linear Constraint Solver (LINCS) (85), and water geometry was preserved using the SETTLE algorithm (86). A timestep of 2 fs was used for all ATMD simulations.

Simulation post-processing was performed using GROMACS (80). Lipid density analysis of coarse-grained simulations was performed with PLUMED (87). A rotxy+transxy fit was applied to remove rotation and translational drift in the xy plane. Lipid contact analysis of atomistic simulations was performed with MDAnalysis (88), with a cut-off of 4 Å. Visualisation used PyMOL (58).

### Supplemental Tables

#### Table S1: Co-evolving residues predicted between YbbA, YbbP and TesA.

**Top co-evolving residues identified between YbbP and TesA**

| Rank | YbbP | TesA | CCMPRED Raw score | Z-score | †Distance (Å) |
|---|---|---|---|---|---|
| 1 | Asp148 | Arg141 | 0.5 | **34.72** | 6.4 |
| 2 | Asp145 | Arg142 | 0.27 | **17.02** | 10.2 |
| 3 | Glu500 | Ala78 | 0.24 | **14.25** | 9.9 |
| 4 | Gln188 | Ala137 | 0.22 | **12.77** | 6.6 |
| 5 | Thr657 | Gln74 | 0.2 | **11.62** | 7.8 |
| 6 | Ala498 | Ala78 | 0.2 | **11.01** | 8.9 |
| 7 | Gln598 | Gln86 | 0.19 | **10.55** | 8.9 |
| 8 | Gly630 | Lys85 | 0.19 | **10.16** | 12.8 |
| 9 | Ala586 | Asp159 | 0.18 | 9.82 | 53.7 |
| 10 | Thr499 | Ala78 | 0.18 | **9.37** | 9.6 |
| 11 | Asp148 | Glu145 | 0.17 | **8.66** | 11.9 |
| 12 | Pro632 | Ile68 | 0.16 | **8.46** | 10.4 |
| 13 | Phe89 | Arg142 | 0.16 | **8.15** | 9.2 |
| 14 | Thr657 | Phe105 | 0.16 | **8.04** | 7.7 |
| 15 | Gln501 | Gln74 | 0.15 | **7.60** | 10.5 |
| 16 | Thr657 | Gln109 | 0.15 | **7.39** | 9.6 |
| 17 | Gln631 | Ala82 | 0.14 | **6.69** | 8.6 |

**Top coevolving residues identified between YbbP and YbbA**

| Rank | YbbP | YbbA | CCMPRED Raw score | Z-score | †Distance (Å) |
|---|---|---|---|---|---|
| 1 | **Gly283** | **Lys84** | **0.24** | **15.25** | 7.3 |
| 2 | Gly720 | Lys84 | 0.17 | **9.65** | 6.9 |
| 3 | Thr279 | Phe92 | 0.17 | **9.59** | 8.2 |
| 4 | Leu273 | Ile100 | 0.17 | **9.24** | 7.1 |
| 5 | Gln709 | Met98 | 0.16 | **8.97** | 9.1 |
| 6 | Leu368 | Ser50 | 0.16 | **8.68** | 12.8 |
| 7 | Phe243 | Ser47 | 0.15 | 7.43 | 66.2 |
| 8 | Leu607 | His10 | 0.13 | 6.59 | 88.0 |
| 9 | Thr716 | Ala81 | 0.13 | 6.37 | 45.3 |

**Top coevolving residues identified between TesA and YbbA**

| Rank | TesA | YbbA | CCMPRED Raw score | Z-score | †Distance (Å) |
|---|---|---|---|---|---|
| 1 | Gly40 | Asn176 | 0.13 | 5.01 | 105 |
| 2 | Pro89 | Val203 | 0.13 | 4.75 | 127 |
| 3 | Ala188 | Ser50 | 0.12 | 4.65 | 108 |
| 4 | Ser73 | Val155 | 0.12 | 4.44 | 107 |
| 5 | Ala146 | Pro165 | 0.12 | 4.34 | 119 |
| 6 | Leu135 | Lys184 | 0.12 | 4.18 | 123 |
| 7 | Leu84 | Ile201 | 0.12 | 4.11 | 123 |
| 8 | Glu157 | Asp187 | 0.12 | 4.03 | 141 |
| 9 | Trp91 | Ph94 | 0.12 | 3.99 | 119 |
| 10 | Leu130 | Asn162 | 0.12 | 3.97 | 117 |

†Distance is between Cα positions in the YbbAP-TesA cryoEM structure.

**Table S2: CryoEM data and model refinement statistics**

| | YbbAP | YbbAP +ATP | YbbAP-TesA +ATP |
|---|---|---|---|
| **Data collection and processing** | | | |
| EMDB code | EMD-51291 | EMD-51292 | EMD-51293 |
| Magnification | 105,000 | 105,000 | 105,000 |
| Voltage (kV) | 300 | 300 | 300 |
| Electron exposure ($e^-$/Å$^2$) | 50 | 50 | 50 |
| Defocus range (μm) | 800-2,300 | 800-2,300 | 800-2,300 |
| Pixel size (Å) | 0.835 | 0.835 | 0.835 |
| Final number of particles | 94,477 | 52,834 | 29,958 |
| Map Resolution (FSC 0.143) | 4.05 | 3.66 | 4.55 |
| **Model refinement** | | | |
| PDB code | 9GE6 | 9GE7 | 9GE8 |
| Resolution (Å) | 4.05 | 3.66 | 4.55 |
| Sharpened Map B | | | |
| Composition | YbbP, YbbA, YbbA | YbbP, YbbA, YbbA | YbbP, YbbA, YbbA, TesA |
| Protein Residues | 799, 224, 224 | 804, 226, 226 | 804, 225, 225, 172 |
| Ligands | - | 2 ANP, 2 $Mg^{2+}$ | 2 ANP, 2 $Mg^{2+}$ |
| B factors | | | |
| Protein | 202, 190, 185 | 168, 138, 131 | 232, 251, 251, 347 |
| Ligands | - | 126, 117, 120, 96 | 237, 237, 209, 228 |
| Rms Deviations | | | |
| RmsBonds | 0.004 | 0.005 | 0.002 |
| RmsAngles | 0.653 | 0.642 | 0.458 |
| **Validation** | | | |
| Map CC (Proteins) | 0.77, 0.84, 0.84 | 0.83, 0.87, 0.87 | 0.79, 0.86, 0.82, 0.82 |
| Map CC (Ligands) | - | 0.94, 0.95, 0.95, 0.97 | 0.89, 0.89, 0.93, 0.95 |
| Q-score | 0.26, 0.31, 0.29 | 0.41, 0.46, 0.46 | 0.26, 0.27, 0.24, 0.19 |
| Clash-score | 19.98 | 13.4 | 10.28 |
| MolProbity score | 2.04 | 2.33 | 1.75 |
| Ramachandran outliers (%) | 0 | 0 | 0 |

**Table S3: Antibiotic susceptibility testing for TesA and YbbP knockout strains.**

| *E. coli* Strain | Plasmid | Antibiotic tested | Notes | MIC Values (µg/mL) | Median MIC (µg/mL) |
|---|---|---|---|---|---|
| BW25113 | - | Penicillin G | - | 32, 32, 32, 32 | 32 |
| *Δtesa* | - | Penicillin G | - | 32, 32 | 32 |
| *Δybbp* | - | Penicillin G | - | 32, 32, 32, 16 | 32 |
| BW25113 | - | Cephalexin | - | 16, 16, 16, 32 | 16 |
| *Δtesa* | - | Cephalexin | - | 16, 16 | 16 |
| *Δybbp* | - | Cephalexin | - | 16, 16, 16, 16 | 16 |
| BW25113 | - | Cephalothin | - | 8, 16 | 12 |
| *Δtesa* | - | Cephalothin | - | 16, 16 | 16 |
| *Δybbp* | - | Cephalothin | - | 16, 16 | 16 |
| C43(DE3) | pET28/Empty | Ampicillin | +IPTG +Kan | 8, 8, 8, 8 | 8 |
| C43(DE3) | pET28/YbbAPTesA | Ampicillin | +IPTG +Kan | 8, 8, 8, 8 | 8 |
| C43(DE3) | pET28/YbbAPTesA(S36A) | Ampicillin | +IPTG +Kan | 4, 4, 4, 8 | 4 |
| C43(DE3) | pET28/YbbAP | Ampicillin | +IPTG +Kan | 4, 4, 8, 8 | 6 |
| C43(DE3) | pET28/Empty | Cephalexin | +IPTG +Kan | 8, 8, 8, 8 | 8 |
| C43(DE3) | pET28/YbbAPTesA(WT) | Cephalexin | +IPTG +Kan | 8, 8, 16, 16 | 12 |
| C43(DE3) | pET28/YbbAPTesA(S36A) | Cephalexin | +IPTG +Kan | 8, 16, 16, 16 | 16 |
| C43(DE3) | pET28/YbbAP | Cephalexin | +IPTG +Kan | 8, 8, 8, 8 | 8 |

All Minimum Inhibitory Concentrations (MICs) measured at 37 °C after 18h – except C43(DE3) experiments which were measured after 48h at 37 °C.

### Supplemental Figures

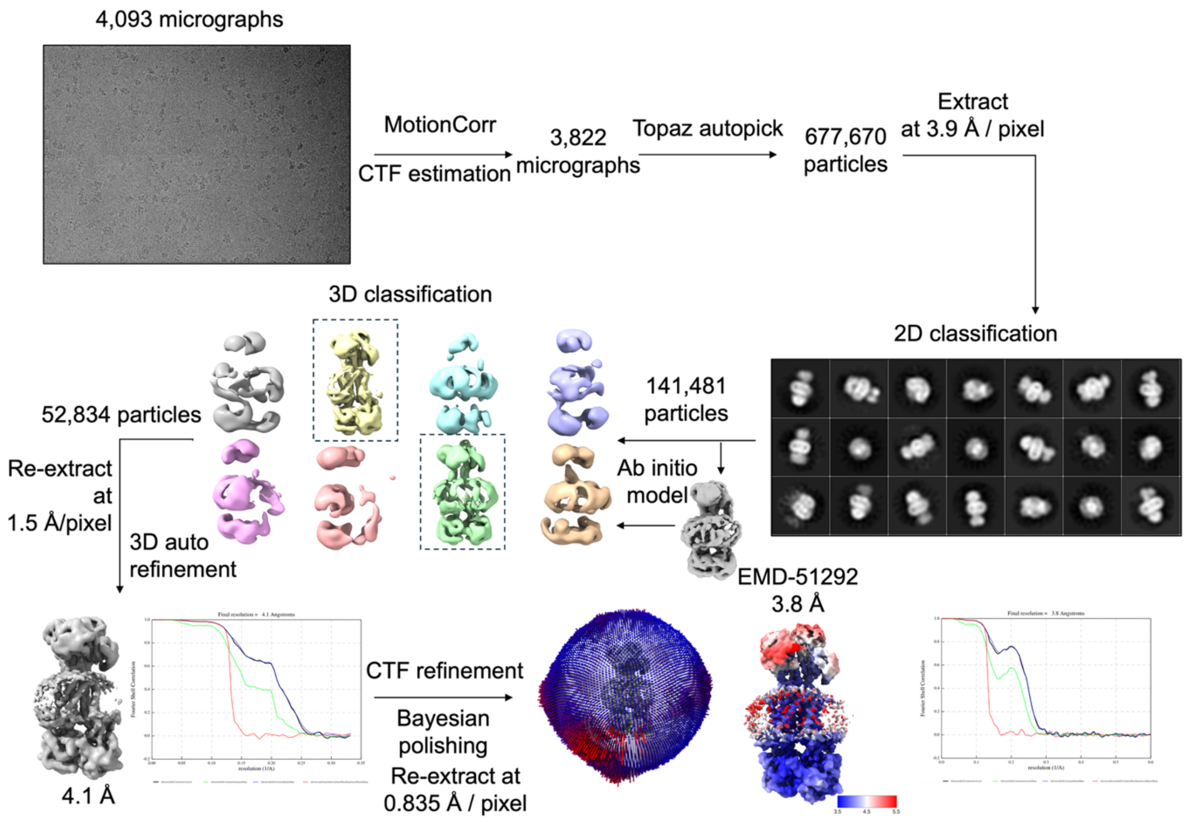


**Figure S1: Image processing workflow for the AMP-PNP bound YbbAP cryo-EM reconstruction.** All processing was performed using Relion with intermediary maps being shown as representatives of multiple jobs. 2D and 3D classifications were performed exhaustively to remove bad particles. The structure is deposited in the Protein Data Bank with accession code 9GE7 with cryoEM maps (sharpened and unsharpened) available from the Electron Microscopy Data Bank under code EMD-51292.

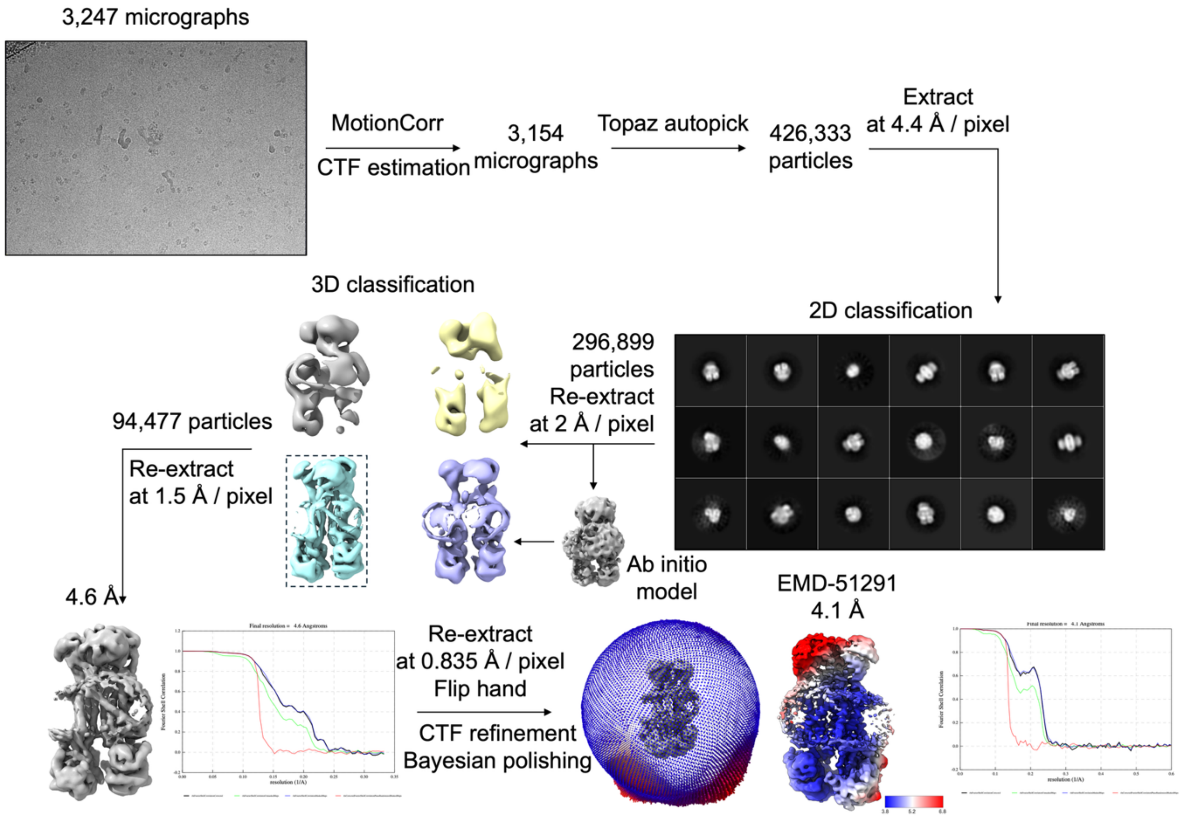


**Figure S2: Image processing workflow for the apo YbbAP cryo-EM reconstruction.** The structure is deposited in the protein data bank with accession code 9GE6. The final maps (sharpened and unsharpened) are available in the EMDB under code EMDB-51291.

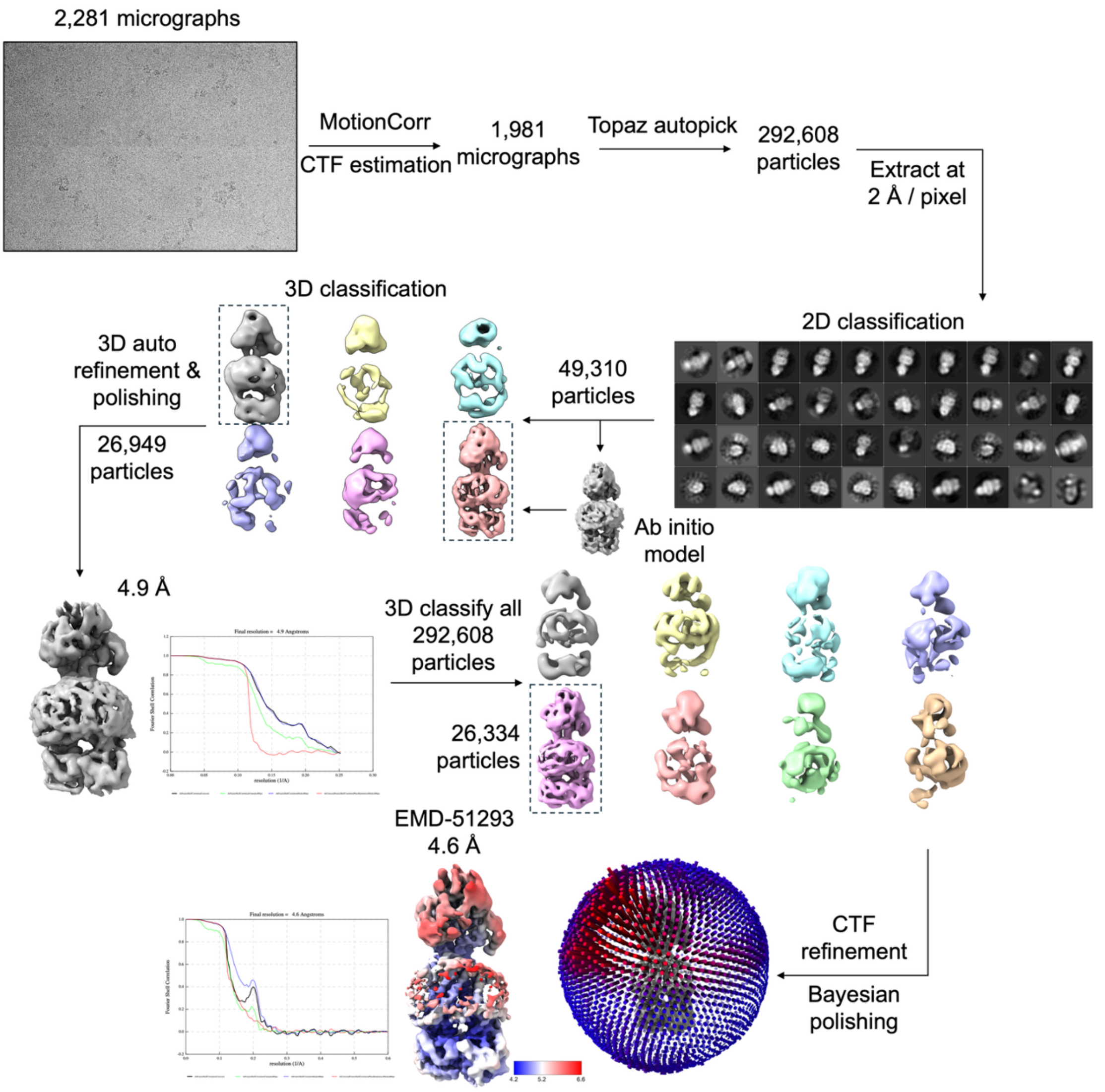


**Figure S3: Image processing workflow for the AMP-PNP bound YbbAP-TesA complex cryo-EM reconstruction**. The structure is deposited in the protein data bank with accession code 9GE8. The cryoEM maps (sharpened and unsharpened) are available in the EMDB under code EMD-51293.

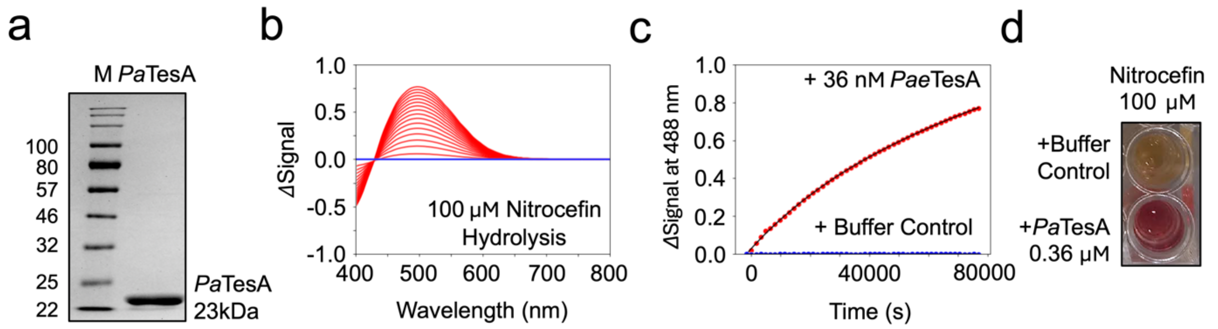


**Figure S4: Nitrocefin and p-nitrophenyl butyrate hydrolysis assays for *P. aeruginosa* TesA**. (a) Purified *Pseudomonas aeruginosa* TesA with N-terminal tag (PaTesA). (b) Representative example of nitrocefin hydrolysis by PaTesA. Red nested curves represent a series of spectral measurements taken over time. Blue curves indicate the control where the enzyme was substituted for protein-free buffer. (c) Plots of the change in absorption at 488 nm for enzyme-catalysed reaction (*red* dots) and the control (*blue* dots). A fit from a single exponential function is shown in black. (d) Photograph of two wells after ~18h showing the colour change associated with nitrocefin hydrolysis.

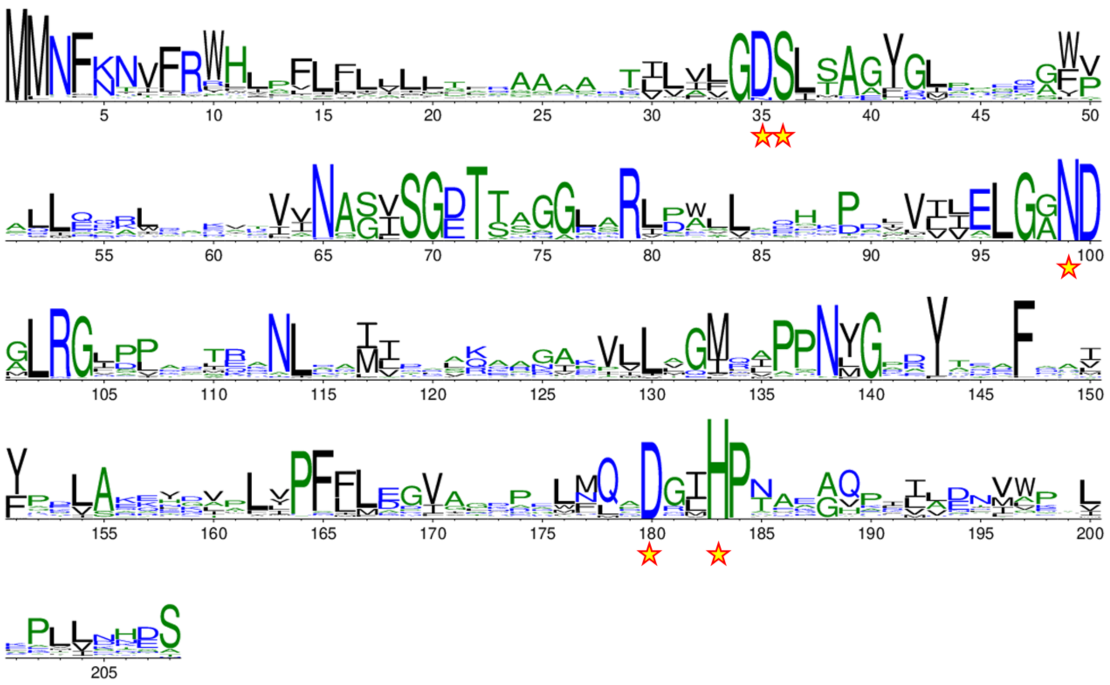

**Figure S5: Amino acid sequence conservation in TesA.** Conservation WebLogo was constructed from a multiple sequence alignment of 12,172 TesA homologues. Stars indicate the positions of key residues within the active site that were tested by mutagenesis (Asp35, Ser36, Asn99, Asp180 and His183).

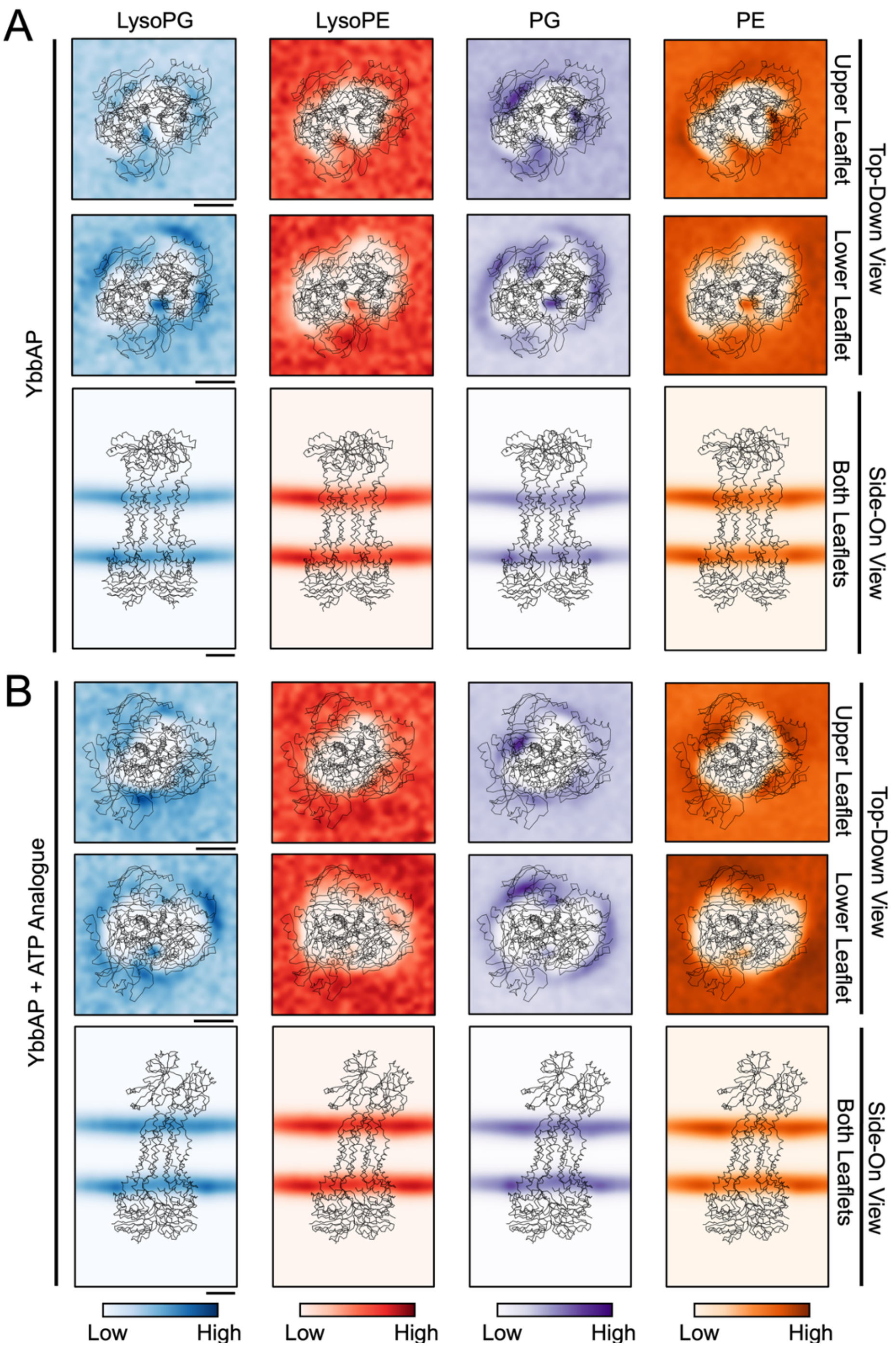


**Figure S6: Lipid residency distributions during coarse-grained simulations of YbbAP.** 2D histograms (bins: 30 x 30) of frequency density of lipid headgroup position (LysoPG - *blue*, LysoPE - *red*, PG – *purple*, PE – *orange*) in the membrane over 15 μs coarse-grained simulation repeats of YbbAP in the (a) apo (n = 3) and (b) nucleotide-bound state (n = 3), normalised to the maximum residency per plot. Calculated from a trajectory of frames comprising of every 10th 1 ns simulation time point. Protein backbone coordinates are shown in *black*. Each scale bar represents 20 Å.

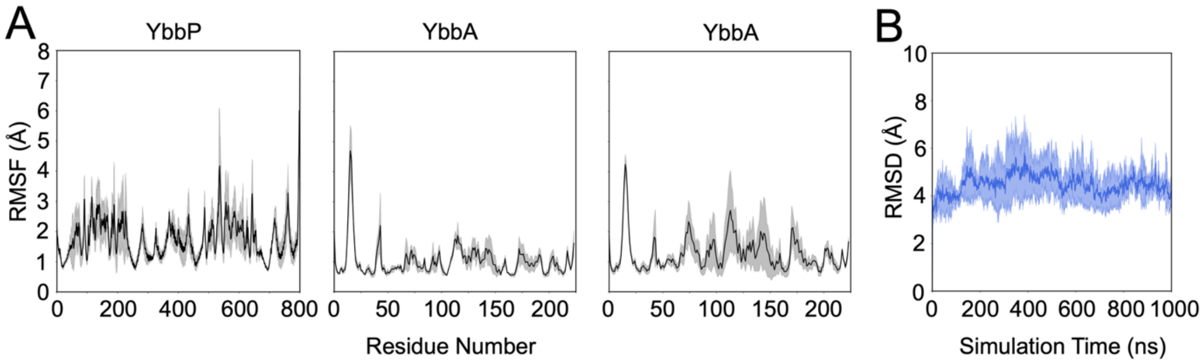


**Figure S7: RMSF and RMSD calculations for atomistic simulations of YbbAP.** (a) Root-mean-square fluctuation (RMSF) per residue over 1000 ns of atomistic simulation repeats (n = 3). Mean RMSF value per residue is shown in *black*, with mean ± 1 standard deviation shown in *grey*. (b) Root-mean-square deviation (RMSD) of YbbAP backbone (in comparison to the start frame) over 1000 ns of atomistic simulation repeats (n = 3). Mean RMSF value per residue is shown in *blue*, mean ± 1 standard deviation shown in *light blue*.

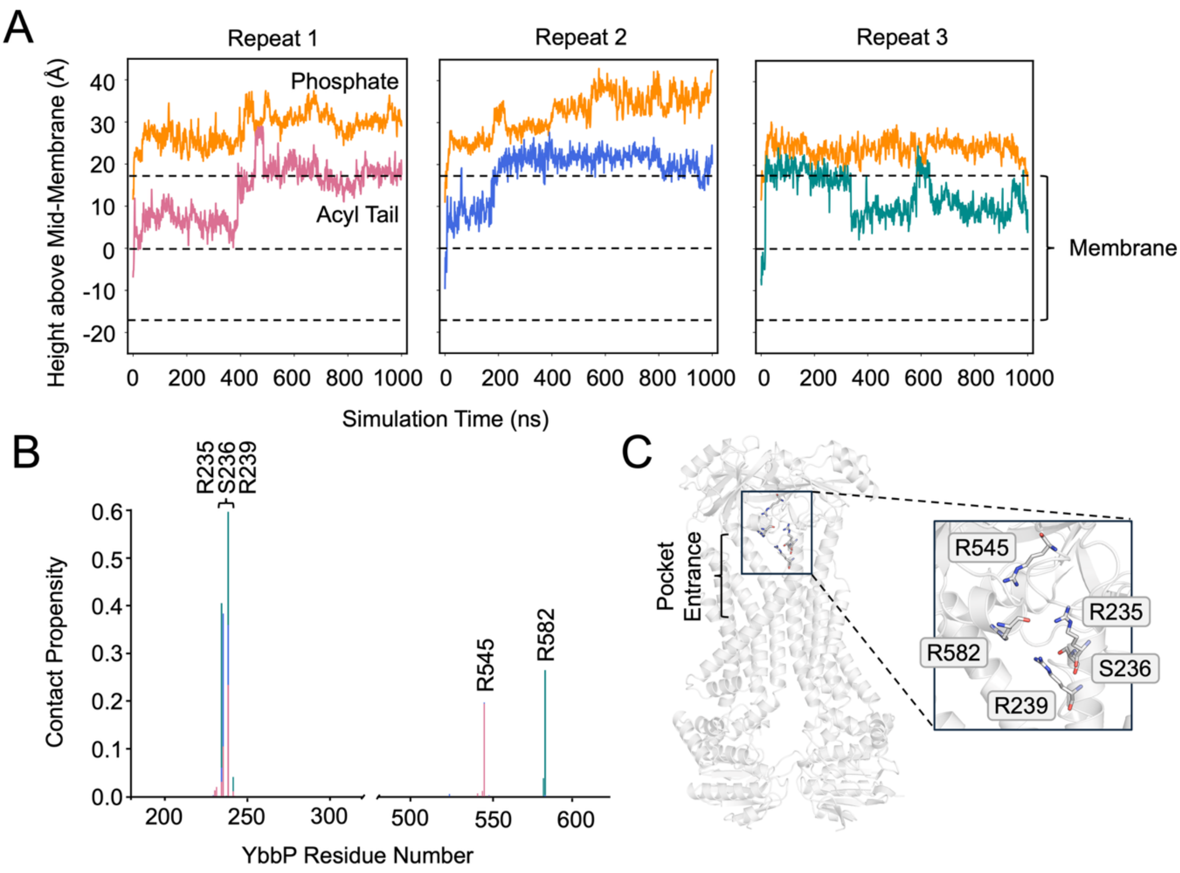


**Figure S8: LysoPG trajectory in atomistic simulation of YbbAP.** (a) Vertical movement of the LysoPG molecule that begins each atomistic simulation repeat within the periplasmic pocket. Repeats 1 (*pink*) and 2 (*blue*) show initial extraction of the phosphate head group, followed by the lipid tail over the course of the simulation. Repeat 3 (*teal*) shows partial extraction of LysoPG from the membrane, which is reversed over the course of the simulation. (b) Contact propensity between the LysoPG and YbbP. Contact Propensity is defined as the fraction of frames where LysoPG is within 4 Å of a given residue. The bar chart shows stacks of values over three repeats. (c) Key residues lining the periplasmic pocket identified during lipid contact analysis that may form interactions with LysoPG during extraction from the bilayer.

### Movie Legends

**Movie 1: Evidence for a mechanotransmission mechanism in YbbAP-TesA.** Molecular morph between ATP bound and nucleotide-free YbbAP structures. (a) View of YbbAP from within the plane of the membrane. (b) View of the cytoplasmic domain movements from within the cytoplasm. Morph generated by Pymol using the cryoEM structures (9GE6 and 9GE7) as start and end points.
